# The temporal balance between self-renewal and differentiation of human neural stem cells requires the Amyloid Precursor Protein

**DOI:** 10.1101/2021.02.17.431707

**Authors:** Khadijeh Shabani, Julien Pigeon, Marwan Benaissa Touil Zariouh, Tengyuan Liu, Azadeh Saffarian, Jun Komatsu, Elise Liu, Natasha Danda, Ridha Limame, Delphine Bohl, Carlos Parras, Bassem A. Hassan

## Abstract

The approximately 16 billion neurons of the human neocortex are derived from a relatively limited number of developmental neural stem cells (NSCs). During embryogenesis, human cortical NSCs initially generate neurons at a particularly slow rate while preserving their progenitor state for a relatively long time. How this balance between the progenitor state and neurogenic state is regulated, and whether it contributes to species-specific brain patterning, is poorly understood. Here we show that the characteristic potential of human NSCs to remain in a progenitor state as they generate neurons for a prolonged amount of time requires the Amyloid Precursor Protein (APP). In contrast, APP is dispensable in mouse NSCs, which undergo neurogenesis at a much faster rate. Mechanistically, loss of APP cell-autonomously accelerates neurogenesis through activation of the AP1 transcription factor and repression of WNT signaling. We propose that the fine balance between self-renewal and differentiation is homeostatically regulated by APP, which may contribute to human-specific temporal patterns of neurogenesis.

## Introduction

The human brain has expanded over fifteen-fold since our divergence from old world monkeys and three-fold since our divergence from a common ancestor with chimpanzees^1^. The cerebral cortex (neocortex) accounts for 2/3 of brain size with enormous diversity in cell type, morphology, connectivity, and function. It is typically structured into six radial layers of cells with highly elaborate neuronal circuits and is responsible for complex cognitive behaviors including sensory perception and skilled motor planning, attention, language, emotion, and consciousness^2,3^. Human cortical neurogenesis begins at around 5 gestational weeks (GW) and is mostly completed by 28 GW^4,5^. An important factor underlying human cortical expansion is an intrinsic capacity of human cortical progenitors to generate very large numbers of neurons over extended periods of time relative to other mammals while retaining a self-renewing progenitor state for a particularly long time, presumably through species-specific mechanisms that remain poorly understood^6,7^.

The highly conserved class I transmembrane Amyloid precursor protein (APP) is broadly expressed throughout nervous system development and has been extensively studied because its proteolytic processing is linked to Alzheimer’s disease AD^8–10^, yet its physiological function, especially in humans, is unclear. APP and its homologues in various species are involved in many biological processes such as axonal outgrowth after injury^11^, endo-lysosomal pathway^12–14^, stress response after hypoxia/ischemia^15,16^, cell signaling processes, and brain development and plasticity^17,18^. In the worm, *C. elegans* loss of the APP homologue *Apl* is lethal due to molting deficits, while causing mild and low penetrance defects in various aspects of neuronal differentiation, function, and survival in mouse and *Drosophila*^19^. In humans, point mutations of even a single copy of *APP* cause familial AD^20^. APP is highly expressed in human telencephalic neurospheres, and during the differentiation and migration of cortical neurons^21–23^ suggesting that it may be involved in NPC proliferation, differentiation and/or maturation^24,25^. In the adult rodent brain, APP is also abundantly expressed in NPCs found in the VZ-SVZ^26,27^, but its loss does not cause obvious deficits in neurogenesis. We generated *APP*-knockout human iPSCs with two different genetic backgrounds and queried the potential effect on human cortical neurogenesis.

## Results

### Loss of APP accelerates cortical differentiation

To examine *APP* expression during human fetal cortical development we queried single cell RNA sequencing (scRNAseq) data from the developing human cortex at GW6-10^28^ (https://cells-test.gi.ucsc.edu/?ds=early-brain) and found that *APP* is present in all six types of cells, with expression in 23.50% of neuroepithelial Cell (NEC), 35.45% of Radial Glial Cell (RGC), 43.66% of Intermediate Progenitor Cell (IPC) and 43.48% of neurons (Fig. 1a-d) present in that dataset. Furthermore, this expression appears particularly dynamic as cells transition from NECs to RGCs and from RGCs to IPCs. We next queried a second scRNAseq dataset from the human developing cortex at GW17-18^22^ and found that APP is expressed in all 16 types of cells present in the human developing cortex (Fig. 1e,f). These data suggest that APP is expressed in human cortical progenitors throughout neurogenesis raising the question of what its function may be at that stage of human cortex development. To gain initial insight into whether APP is potentially required during human fetal development, we queried genetic loss of function data obtained from the human population by the GNOMAD project^29,30^. These analyses reveal that APP has a loss of function observed/expected (LOEUF) score of 0.42 (https://gnomad.broadinstitute.org/gene/ENSG00000142192?dataset=gnomad_r2_1), meaning that complete loss of APP causes ∼60% developmental mortality, a rate surprisingly higher than that of key neurodevelopmental genes such as Neurog2 (LOEUF=0.54) and Ascl1 (LOEUF=0.59) which show less than 50% developmental mortality (Fig. 1g). Together, these data suggest that APP may play an important role in human cortex development.

**Fig. 1.**
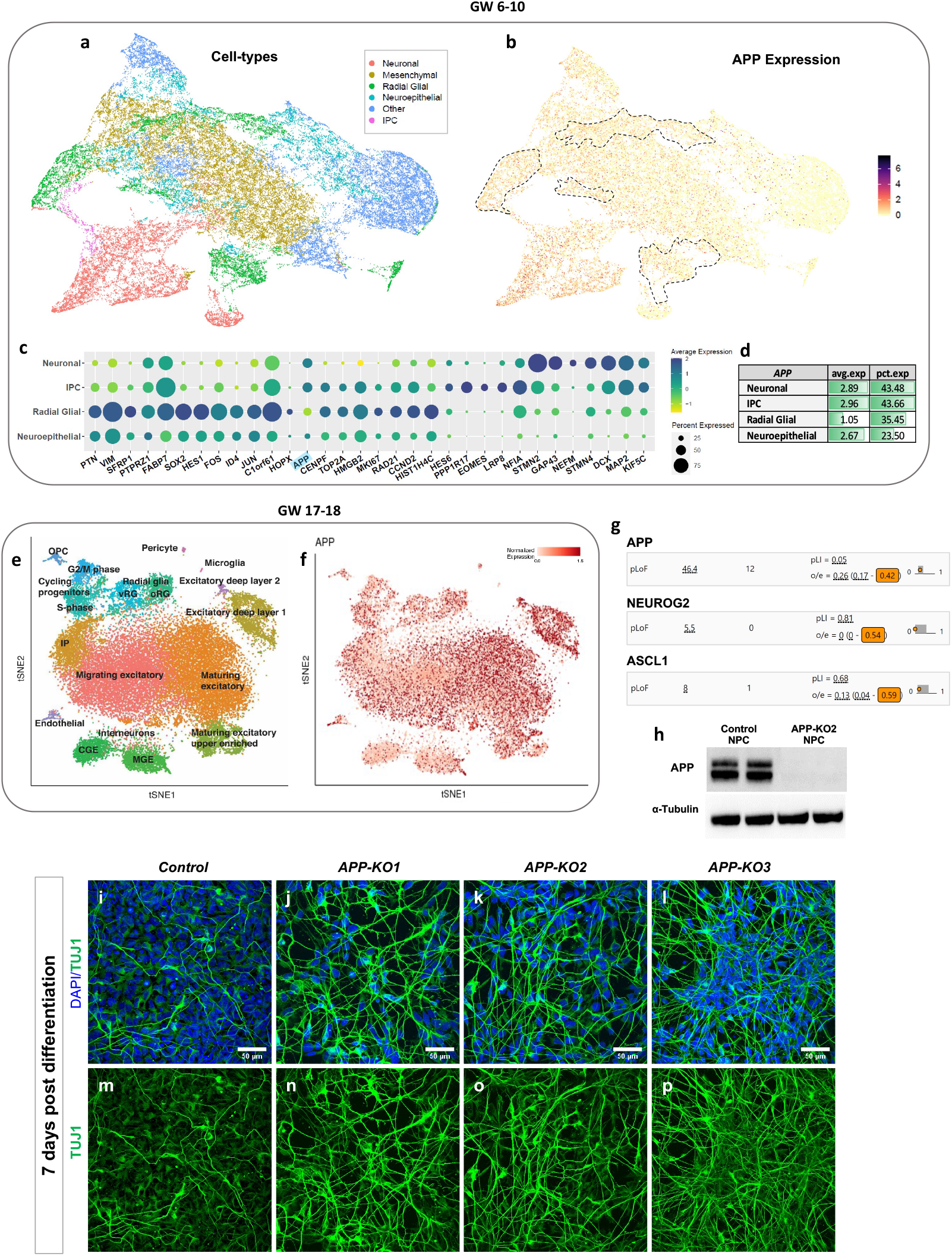
APP expression in the developing human cortex. **a-d**, APP is expressed in all cell types detected by scRNAseq of the human fetal cortex during GW6-10 and **e-f**, GW17-18 of development. **g**, *APP* loss of function results in significant developmental lethality in the human population with a lower observed/expected (LOEUF) score (0.42) compared to two key neurodevelopmental genes such as NEUROG2 (LOEUF=0.54) and ASCL1 (LOEUF=0.59). **h**, APP expression in control iPSC-derived human NPCs was confirmed by western blot. **i-p**, Many more cells express the neuronal markers TUJ1 in in all *APP-KOs* compared to isogenic control 7 days post differentiation (Scale bar 50 µm).

To test this hypothesis, we generated *APP-KO* iPSCs by CRISPR/Cas9 (Extended Data Fig. 1a-k, see methods for iPSC lines and experimental details) and chose three *APP-KO* clones and one *APP* control clone, which received Cas9 but was not mutated, for further analysis. We verified the karyotype and the APP locus in all clones to ensure isogenicity. Upon induction of cortical differentiation, the morphology and molecular identity of NPCs were normal in all clones (Extended Data Fig. 2). We first confirmed that APP protein is expressed in human cortical progenitors (Fig. 1h). Seven days after initiation of neuronal differentiation, we stained the cells for SOX2 (NPCs), Doublecortin (DCX; immature neurons), β-TubulinIII (TUJ1; neurons) to determine cell type. We observed a striking increase in TUJ1 (Fig. 1i-p) and DCX (Extended Data Fig. 3a-h) expression 7 days post differentiation in all three *APP-KO* clones relative to the control clone. In order to examine whether this effect is general or brain-region specific, we generated motor neurons^31^ from control and *APP-KO* iPSCs. While human cortical neurogenesis occurs over 3-4 months, spinal motor neurons are generated at a much faster rate in 2-3 weeks. We used a 17-days protocol with 10 days of neural induction and 7 days of neural differentiation. We chose two early time points (day 1 and day 4 post-differentiation; i.e. days 11 and 14 of culture) which approximately correspond to the 7 days post-differentiation time point in cortical neurons (Extended Data Fig. 4a). We first confirmed the expression of APP protein in motor neuron progenitor cells (day 10 of culture; Extended Data Fig. 4b) then examined their differentiating daughters for expression of ISLET1 (MN marker) and TUJ1 (Extended Data Fig. 4c-n). No significant differences were detected in the number of ISLET1+ cells in *APP-KO2* and control (Extended Data Fig. 4o) at either time point. These data suggest that loss of *APP* preferentially affects the slower cortical progenitors compared to the faster motor precursors.

To determine the neurogenic potential of control and *APP-KO* cortical NPCs, we sparsely labeled them (70 control NPCs and an average of 63.6 NPCs in the 3 *APP-KO* clones, Supplementary Table 1) with a GFP-expressing lentiviral vector^32,33^ (Fig. 2a) and quantified the number of GFP+ progenitors (SOX2) and neurons (TUJ1) at days 0, 7, and 30 after differentiation (Fig. 2b-u). After 7 days, we observed a significant decrease in NPCs (Fig. 2v) paralleled by a significant increase in neurons over time (Fig. 2w), in all three *APP-KO* compared to control. Of note, while the proportion of neurons generated by *APP-KO* progenitors shows a plateau between days 7 and 30, the slope is still rising in controls (arrow in Fig. 2w) meaning that neurogenesis is still ongoing in the control background, consistent with previous reports^33^. Quantitatively, we found that 1 control NPC produced on average 8 NPCs and 3.6 neurons after 7 days of differentiation. In contrast, within the same time frame, each *APP-KO* NPC produced on average 1.25 NPCs and 22.9 neurons (Supplementary Table 1), indicating a major switch of cortical NPCs from self-amplifying state to neurogenic state.

**Fig. 2.**
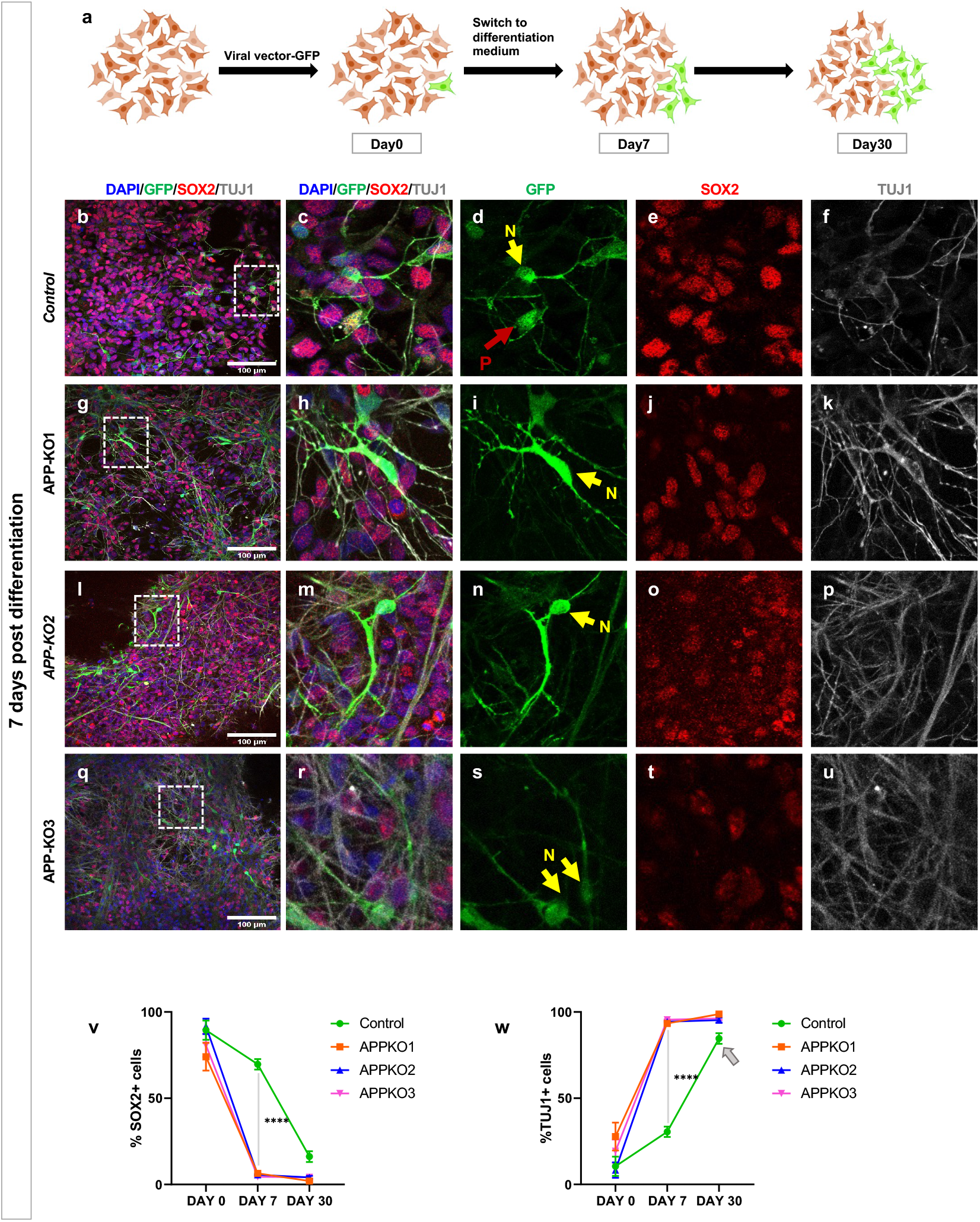
Following neurogenesis using sparse labeling of progenitors suggests accelerated neurogenesis in APP-KO clones. **a**, Schematic illustration of the sparse labelling approach using a lentiviral vector to follow the fate of the progeny of labelled progenitors (icons adapted from BioRender.com). **b-u**, Expression of SOX2 and TUJ1 within GFP+ cells in isogenic control and *APP-KOs* 7 days post differentiation (Scale bar 100 µm, N=neuron, P=Progenitor). **v**, Decrease in SOX2 and **w**, Increase in TUJ1+ expression in *APP-KO* clones compared to isogenic control within 7 days (2way ANOVA, p<0.0001). At 30 days post-differentiation the ratios of progenitors and neurons have plateaued in *APP-KO* clones. The ratios in isogenic control clone have reached almost similar levels as in KO clones but they are still on a dropping (progenitors) and rising (neurons, grey arrow) trajectories, respectively.

### Cortical neurons are generated in a shorter time window in the *APP-KO* background

One of the key features of cortical development is the sequential generation of different neuronal subtypes that will occupy different layers of the neocortex. This feature is preserved in NPC cultures, whereby early-born (deep layer) neuronal fate markers appear before late-born (upper layer) cell fate markers^7,34^. To assess whether loss of APP affects this order, differentiation was launched and cells were tracked for 7, 15, 30, and 50 days post-differentiation. We first observed clear expression of early-born neuronal marker CTIP2 only at day 15 in both genotypes, despite the fact that *APP-KO* NPCs had generated ∼3-fold more neurons than control NPCs (Extended Data Fig. 5a1-f4). Within 30 days, the number of CTIP2+ and STAB2+ cells increased in both control and *APP-KO2* (Extended Data Fig. 5g1-h4). The same results were observed for *APP-KO1* and *APP-KO3* at 7, 15, and 30 days post-differentiation (data not shown). Thus, *APP-KO* progenitors are capable of generating early-born and late-born neurons in the correct sequence.

Put together, our data so far can be explained by two very different models. In a first model, which we term the “interrupted neurogenesis” model, *APP-KO* progenitors initiate neurogenesis prematurely causing an exhaustion of the progenitor pool while they are still in their early temporal phases. In a second model, which we term the “accelerated neurogenesis” model, *APP-KO* progenitors go through their entire temporal program but in a much shorter period. These two models make diametrically opposite predictions for the early born to late born neuron ratio (EB/LB ratio) at an intermediate stage of neurogenesis. The first model predicts that the EB/LB ratio would be higher in *APP-KO* compared to control (Fig. 3a) because the system would run out of progenitors before enough late born neurons are produced. The second model predicts that the EB/LB ratio would be lower in *APP-KO* compared to control (Fig. 3b) because APP-KO progenitors would have completed their temporal program while control progenitors are still in early phases of generating late born neurons. To test these predictions, we stained 50 days post-differentiation cultures for the neuronal fate markers FOXP2 (EB) and SATB2 (LB) and found that the ratio of FOXP2/SATB2 (EB/LB ratio) is consistently lower in all 3 *APP-KO* clones compared to the control reaching significance in 2 of the 3 (Fig. 3c-o). These data are consistent with model 2 and suggest that APP contributes specifically to the absolute temporal scale of human neurogenesis, but not the relative temporal birth order of cortical neurons.

**Fig. 3.**
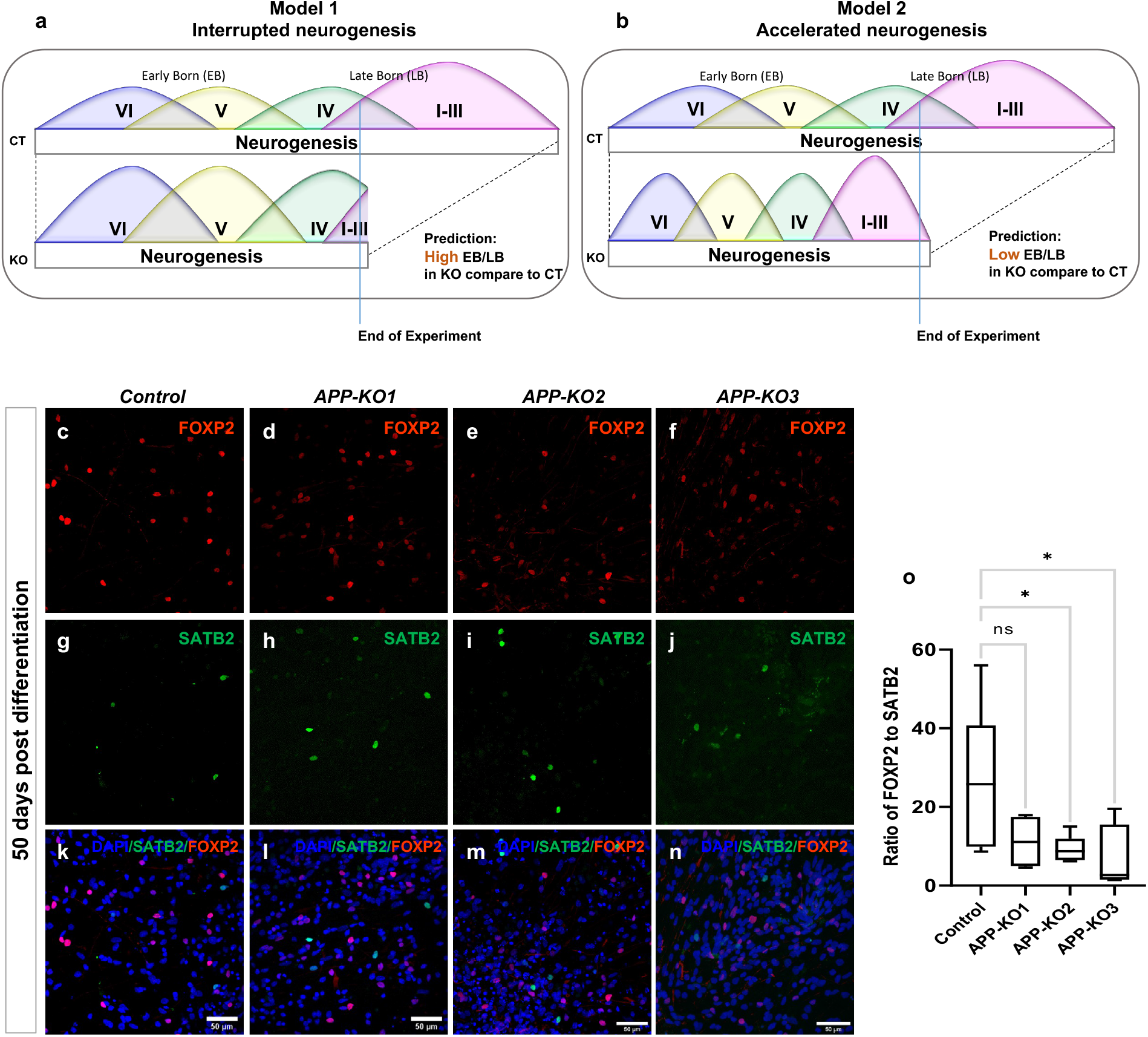
Accelerated production of all cortical neuron types in *APP-KO* clones. **a**, The “Interrupted neurogenesis” model in which APP KO progenitors are prematurely depleted before producing sufficient late born neurons predicts a high early born (EB) to late born (LB) neuron ratio compared to isogenic controls **b**, Conversely, the “Accelerated neurogenesis” model in which APP KO progenitors undergo the same temporal neurogenic series but with faster dynamics predicts a low EB/LB neuron ratio at the same time point during active neurogenesis. **c-n**, 50 days post differentiation neurons stained for FOXP2 and SATB2 in isogenic control and *APP-KOs* (scale bar 50µm). **o**, Low ratio of EB/LB neurons in APP-KO compared to isogenic control supports the accelerated neurogenesis model in the absence of APP (n=1, Ordinary one-way ANOVA, p=0.0557 for isogenic control vs APP-KO1, p=0.0373 for isogenic control vs APP-KO2, p=0.0252 for isogenic control vs APP-KO3).

### Loss of APP accelerates differentiation of human NPCs

Given the expression of APP in fetal human cortical progenitors^22,28^ (Fig. 1a-f) and the acceleration of neurogenesis observed in *APP-KO* cells, we hypothesized that that APP is required in the progenitors themselves to maintain cortical NPCs in a prolonged self-amplifying progenitor state. To test this idea, we quantified the ratio of NPCs versus IPCs and early born neurons by staining pure NPC cultures for SOX2 and DCX (Fig. 4a-o) prior to the onset of neuronal differentiation, where there should be very little spontaneous differentiation under control conditions. Strikingly, we observed a significant decrease in the ratio of SOX2+ cells (Fig. 4p) and a significant increase in the ratio of DCX+ cells (Fig. 4q) in *APP-KO*s relative to the isogenic parental (see materials and methods) and gRNA/Cas9 treated controls. To rule out genetic background effects, we generated an *APP-KO* clone from a different independent iPSC line (iPSC line 2; see methods for details) (Extended Data Fig. 1l-n) and observed the same results in those NPCs (Extended Data Fig. 6a-k). We also produced cortical organoids from control and *APP-KO* clones of this line and stained them for SOX2 and DCX at day 15, when the majority of cells are in the NPC state (Extended Data Fig. 6l-s and supplementary videos I and II). We performed 3D reconstructions of the organoids and quantified DCX expression. We observed both more and larger DCX+ clusters in *APP-KO* organoids relative to control (Extended Data Fig. 6t,u).

**Fig. 4.**
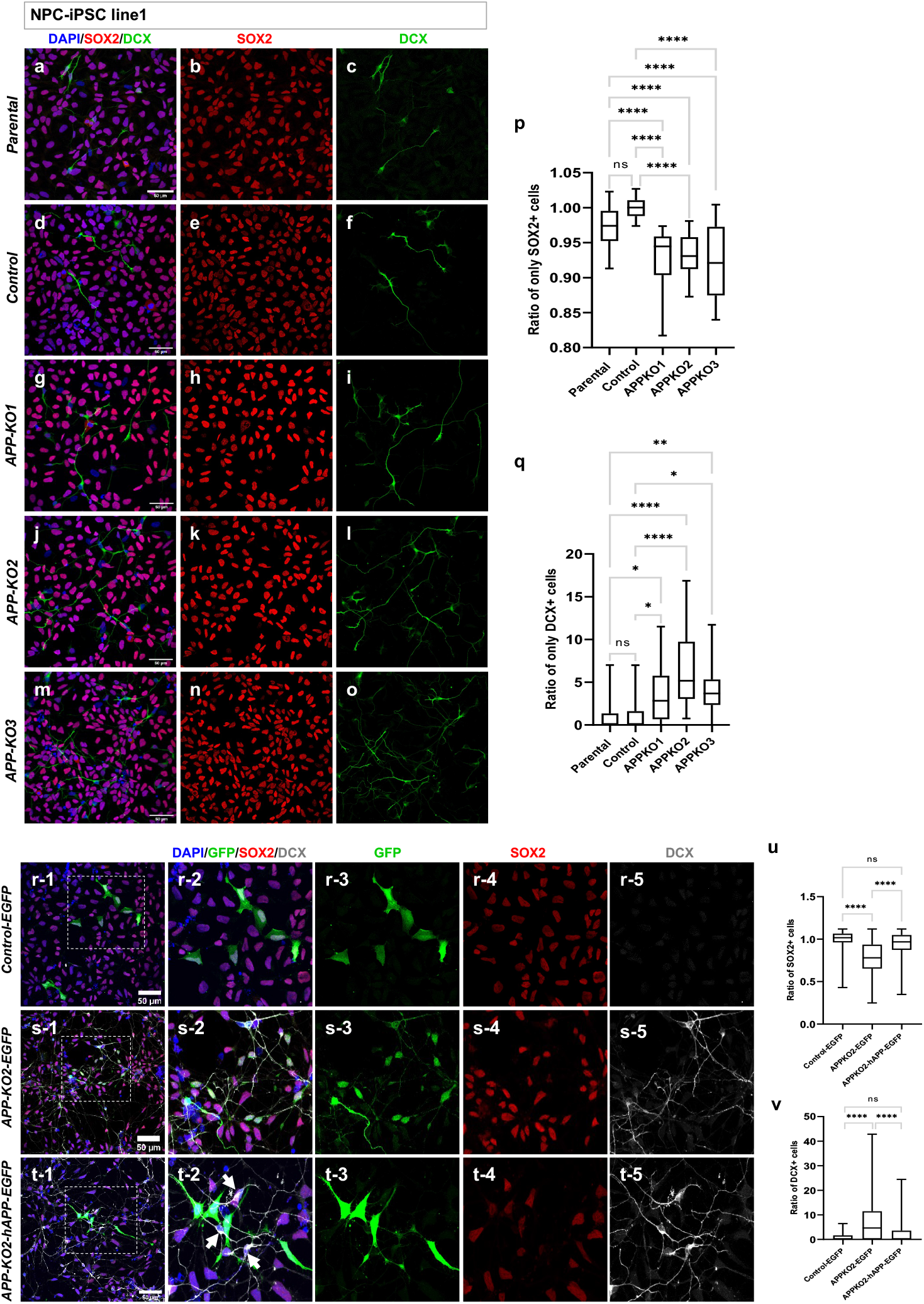
Loss of APP drives premature differentiation of human NPCs. **a-o**, NPCs in proliferating medium from parental, isogenic control and *APP-KOs* stained for SOX2/DCX. **p**, Significant decrease in ratio of SOX2+ cells (n=3, one-way ANOVA, p<0.0001 for all the conditions) and **q**, Significant increase in ratio of DCX+ in *APP-KOs* background compare to parental and isogenic control (n=3, one-way ANOVA, p=0.0235, p<0.0001, p=0.0061 for parental vs APP-KO1, APP-KO2 and APP-KO3 respectively. p=0.0487, p<0.0001, p=0.0151 for isogenic control vs APP-KO1, APP-KO2 and APP-KO3 respectively, scale bar 50µm). **r1-r5**, Isogenic control NPCs transfected with *pPB-CAG-IRES-EGFP* **s1-s5**, *APP-KO2* NPCs transfected with *pPB-CAG-IRES-EGFP* **t1-t5**, *APP-KO2* NPCs transfected with *pPB-CAG-hAPP-IRES-EGFP* and stained for GFP/SOX2/DCX. APP rescued the phenotype of transfected cells but not neighboring cells (arrows in T2). **u**, Ratio of SOX2+ and **v**, DCX+ cells in all 3 conditions (n=3, Ordinary one-way ANOVA, p<0.0001, scale bar 50µm).

Next, we performed a rescue experiment by transfecting *APP-KO2* NPCs with PiggyBac vectors containing either GFP alone (*pPB-CAG-IRES-EGFP*) or GFP and APP (*pPB-CAG-hAPP-IRES-EGFP*) and staining for SOX2 and DCX. Quantification of GFP+/SOX2+ and GFP+/DCX+ cells showed that *APP-KO* NPCs that received the APP containing vector but not GFP alone were restored to a progenitor state and the number of their neuronal progeny was reduced back to control levels (Fig. 4r1-v). We noted that restoring APP only rescued NPCs expressing it, but not neighboring cells (arrows in Fig. 4t2), suggesting a cell-autonomous requirement. Together, these data show that APP is cell-autonomously required to repress premature neurogenesis in human cortical NPCs.

Finally, to test whether this effect on cortical neurogenesis was specific to human cortical NPCs, we tested for signs of premature neuronal differentiation in the *APP-KO* mouse cortex at E10.5, a stage when most cells should still be progenitors. Previous studies of the role of APP and its paralogues APLP1 and APLP2 in mouse neurogenesis have consistently reported contradictory results, including increased neurogenesis, decreased neurogenesis and absence of any effects ^35–38^. We examined the developing mouse cortex at E10.5 and observed no Tuj1+ cells in the progenitor layer and no difference in Pax6 (NPC) and Tuj1 expression between *App-WT* and *App-KO* brains (Extended Data Fig. 7a-n). We then examined the number of progenitors (Pax6), early-born (Ctip2), and late-born (Satb2) neurons at E17.5 towards the end of neurogenesis. We observed no significant change in the number of Satb2, Ctip2, or Pax6 positive cells between *App-WT* and *App-KO* brains (Extended Data Fig. 7o-z’). These results suggest that APP’s requirement in cortical progenitors may not be as critical in the mouse cortex as it is in humans, perhaps because mouse cortical neurogenesis is much faster.

### Loss of APP accelerates the differentiation of human cortical NPCs

To gain insight into how APP regulates the fate of human cortical NPCs, we performed single cell RNA sequencing using a novel, non-microfluidics-based method, on control and *APP-KO* NPCs (Supplementary Table 2, BioProject accession number PRJNA678443). First, we confirmed that both control and *APP-KO* NPCs mapped to progenitor cells in the human fetal brain by integrating them with recently reported single cell transcriptomes^28^ using Integrated Anchors analysis followed by clustering with Seurat^39^ (Fig. 5a,b). Cluster analysis without taking into account cell identity identified seven clusters, six of which showed an appropriate distribution following lineage progression as shown by uniform manifold approximation and projection (UMAP) 2D-representation (Fig. 5c). However, cluster 4 was located apart from the rest and did not fit within the smooth lineage progression of the remaining clusters. This cluster contained both control and *APP-KO* cells, with the majority being *APP-KO* (Fig. 5d), most of which mapped to neural progenitors of human fetal brain cells (Fig. 5e). To more precisely identify these cells, we performed clustering analyses according to cell identity, following the cell-type annotation of Polioudakis et al.^22^, which confirmed the presence of closely related control and *APP-KO* clusters (UMAP 2D-representation; Extended Data Fig. 8a and Heatmaps; Extended Data Fig. 8c,d) of radial glial cells (RGC), cycling progenitors 1 and 2 (CP1 and CP2), intermediate progenitors (IP) and neurons (N), as well as a separate cluster of KO cells that we refer to as the neurogenic progenitors (NP) cluster (see below). These cells represent the majority of KO cells in cluster 4 of the previous analysis. The distribution and percentage of cells per cluster show that NPs represent 13% of the *APP-KO* population and together with the neurons appear at the expense of cycling progenitors, which show a reduction of ∼20% in *APP-KO* cells compared to control (Extended Data Fig. 8b), suggesting premature cell cycle exit and increased differentiation of *APP-KO* cells. Consistent with this, we found a 20% reduction in cells expressing the cell cycle marker Ki67 in *APP-KO* NPCs compared to controls (Extended Data Fig. 8e-g).

**Fig. 5.**
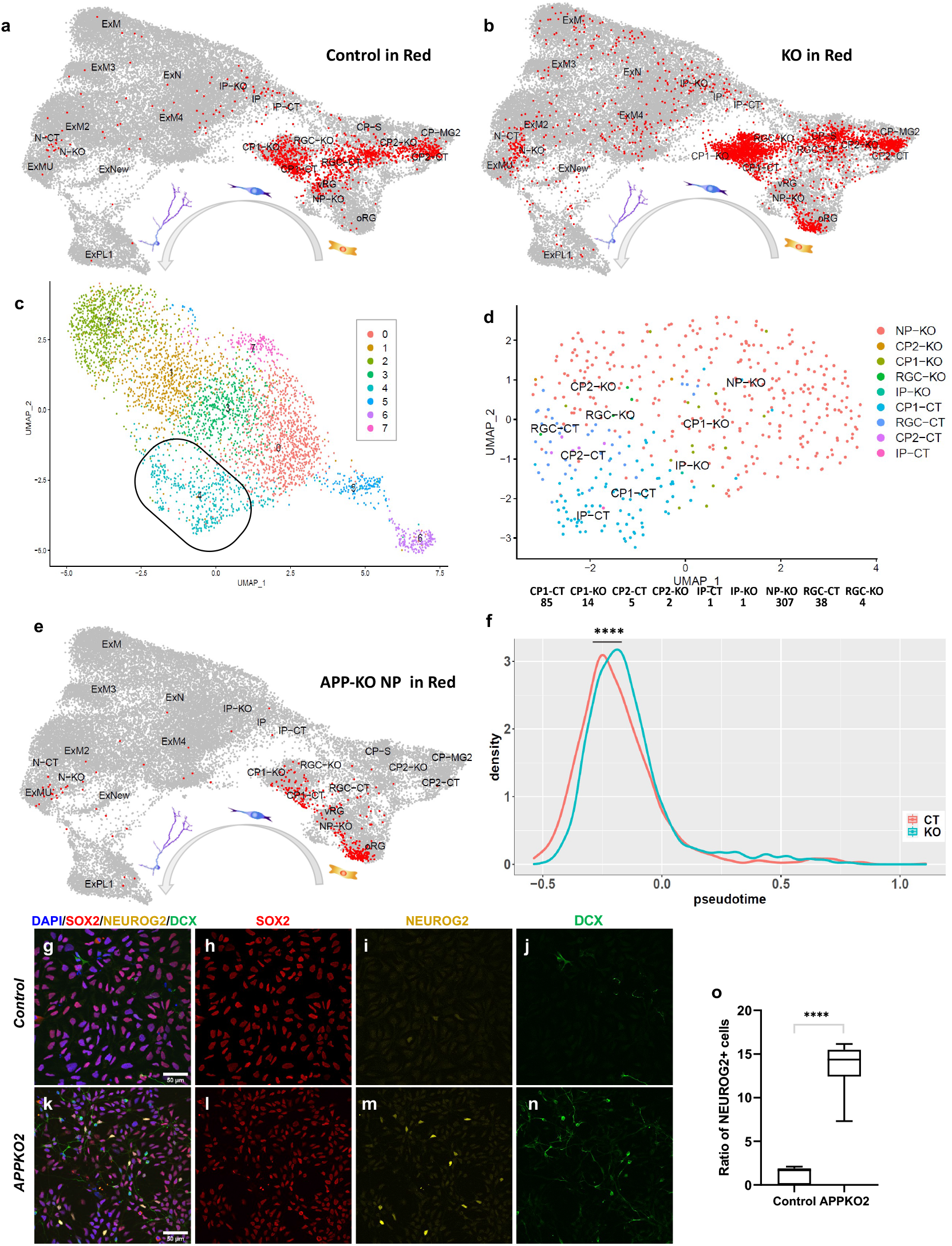
Premature transition to a neurogenic state in *APP-KO* NPCs. **a-b**, sc-RNA sequencing data from isogenic control and APP-KO NPCs both map to normal human progenitors cell types. **c**, Cluster analysis without *a priori* cell identity identified seven clusters most of which are composed of similar ratios of isogenic control and *APP-KO* cells (see Extended Data Fig. 8) with the exception of cluster 4. **d-e**, The majority of cells in cluster 4 are *APP-KO* NPCs that map to normal human fetal progenitors. **f**, Pseudotime analysis of the scRNAseq data shows a temporal shift towards a differentiated state in *APP-KO* NPCs compared to isogenic control (p-value=8.418e-13, Kolmogorov-Smirnov test). **g-n**, Staining of isogenic control and *APP-KO2* NPCs for NEUROG2/SOX2/DCX. **o**, Ratio of NEUROG2+ neurogenic NPCs is 15-fold higher in *APP-KO2* compared to isogenic control (unpaired t-test, p<0.0001, scale bar 50µm).

Our scRNAseq dataset gave us the opportunity to further test the accelerated neurogenesis model by performing pseudotime analysis to determine the temporal trajectory of *APP-KO* NPCs relative to controls. Pseudotime analysis shows a significant overall temporal shift towards a differentiated state in the *APP-KO* population compared to control (Fig. 5f) which was true for all classes of progenitors but not IPs and neurons (Extended Data Fig. 8h). To further complement these observations, we performed SOX2/NEUROG2/DCX triple staining (Fig. 5g-n) on control and *APP-KO* NPCs. NEUROG2 expression marks commitment to differentiation of mammalian NPCs^40,41^. We observed a ∼15-fold increase in the number of NEUROG2+ NPCs in *APP-KO* compared to control (Fig. 5o). This is close to the increased ratio of neurons-to-progenitors (∼18-fold) we found to be produced by *APP-KO* NPCs (Supplementary Table 1). Together, our data show that loss of *APP* creates an accelerated drive towards neurogenesis in human cortical progenitors, suggesting that APP regulates downstream mechanisms required to maintain cortical NPCs in a progenitor state.

### Loss of APP in cortical NPCs triggers AP1 activation to drive neurogenesis

To gain insight into the mechanisms of the gain of neurogenic state in the absence of APP, we performed bulk RNA sequencing in *APP-KO* and control NPCs. We obtained 763 differentially expressed genes (558 up; 205 down including *APP* as expected) between control and *APP-KO2* NPC (Supplementary Table 3 and Extended Data Fig. 9a,b). Consistent with the *APP-KO* phenotype, bulk RNAseq showed downregulation of primary progenitor’s markers *PAX6* and *CDON*, and upregulation of neurogenic genes *NEUROG2, BCL6, ASCL1, NHLH1*, and *DCX*. Moreover, *CDKN2B*, a well-known cell cycle inhibitor, and differentiation promoting factors *DLL3, JAG2*, and *DNER* were also upregulated (Extended Data Fig. 9c). One pathway whose activity is intimately linked to APP function in a variety of organisms and neuronal contexts is the JUN N-terminal Kinase (JNK) pathway^42^ whose effector proteins, JUN and FOS, combine to form the AP1 transcription factor. Significant evidence supports APP-dependent transcriptional repression of c-Jun and reduced basal activity of c-Jun N-terminal kinase (JNK) in PC12 cells^43^ with activation of the JNK signaling pathway reduced either by APP overexpression or treatment with sAPPα^43–45^. Moreover, *c-Fos* expression is upregulated in the prefrontal cortex of *App-KO* mice^46,47^. We observed increased expression of *JUN* and *FOS* mRNAs in *APP-KO* NPCs (1.3 and 2.7 Log2FC, respectively; Fig. 6a), suggesting that APP also regulates their expression in human context. JUN is activated upon phosphorylation by JNK^48–50^, and we observed a 1.5-fold increase in phospho-JUN in *APP-KO* NPCs compared to control (Fig. 6b,c). Finally, AP1 regulates the expression of BCL6 in germinal center B cells^51^, and BCL6, which we found to be upregulated in *APP-KO* NPCs, drives neurogenesis of cortical progenitors in the developing mouse cortex ^52,53^. Analysis of genes characterizing the *APP-KO* NP cluster of our single cell transcriptomic dataset showed enrichment for binding of JUN and JUND to their promoters by chromatin immunoprecipitation (ChIP-seq) (ChEA 2016; Fig. 6d). Furthermore, we also found consensus AP1 binding sites in the regulatory regions of genes upregulated in *APP-KO* NPCs in the bulk RNAseq data, including *DCX* (Supplementary Table 4). To test whether AP1 activation might drive *APP-KO* NPCs into a neurogenic state, we treated control and *APP-KO2* NPCs with a specific AP1 inhibitor (SR11302)^54,55^ and stained them for SOX2, NEUROG2, and DCX (Fig. 6e-t). While the ratio of SOX2+ cells remained unchanged after treatment (Fig. 6u), the number of NEUROG2+ and DCX+ cells was significantly rescued (Fig. 6v,w). These results suggest that loss of APP causes premature differentiation through AP1 activation independently of NPC self-renewal. This indicates a second APP-dependent mechanism for NPC amplification.

**Fig. 6.**
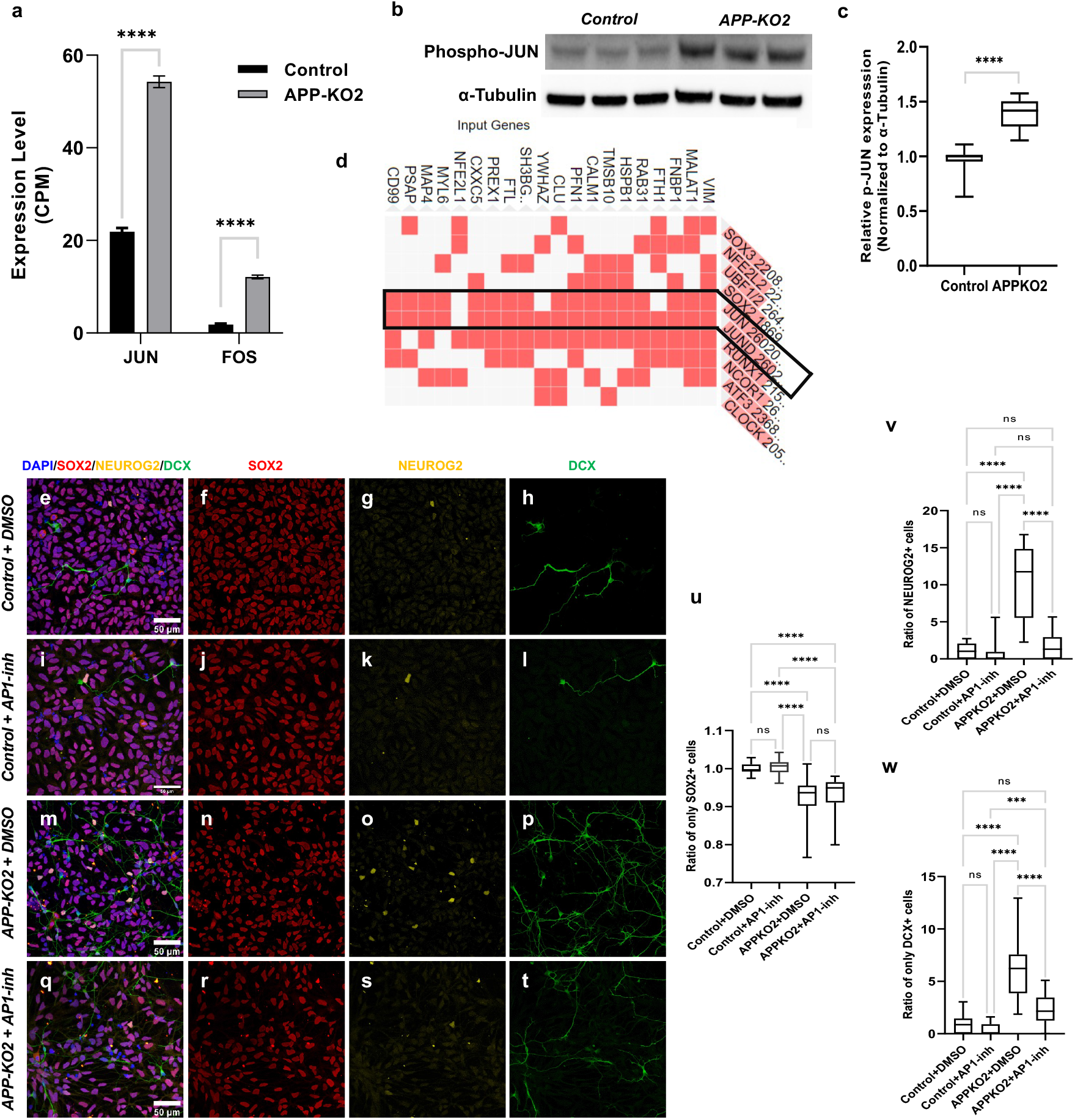
Loss of APP increases differentiation of NPCs through activating the JNK pathway. **a**, Upregulation of *JUN* and *FOS* in *APP-KO2* NPCs in bulk RNA sequencing results (Ordinary one-way ANOVA, p<0.0001). **b-c**, Western blot shows significant increase in phospho-JUN in *APP-KO2* NPCs compared to isogenic control (n=3 biologically independent repeats, unpaired t-test, P<0.0001). **d**, JUN family members bind to marker genes of neurogenic progenitors (NPs) in the ChEA 2016 ChIP-seq database. **e-t**, NPCs treated with AP1 inhibitor (AP1-inh, SR11302) and stained for SOX2/DCX/NEUROG2 showed **u**, No increase in ratio of SOX2+ cells in *APP-KO2* treated with SR11302 compared to untreated *APP-KO2* NPCs (n=3, Ordinary one-way ANOVA, p<0.0001 for all the significant condition). **v**, Significant decrease in NEUROG2+ cells in *APP-KO2* treated with SR11302 compared to untreated *APP-KO2* NPCs (n=3, Ordinary one-way ANOVA, p<0.0001 for all the significant conditions). **w**, Significant decrease in DCX+ cells in *APP-KO2* treated with SR11302 compared to untreated *APP-KO2* NPCs (n=3, Ordinary one-way ANOVA, p<0.0001 for isogenic control-DMSO vs *APP-KO2*-DMSO, isogenic control-SR11302 vs *APP-KO2*-DMSO and *APP-KO2*-DMSO vs *APP-KO2*-SR11302 and p=0.0008 for isogenic control-SR11302 vs *APP-KO2*-SR11302). Scale bar 50µm for all the images.

### Loss of APP reduces NPC amplification by attenuating the canonical WNT pathway

We have recently shown that APP is a receptor for Wnt3a and that Wnt3a binding protects APP from degradation^56^. In addition, KEGG pathway analysis of our bulk RNAseq data indicated that WNT signaling, which is required in a variety of contexts for stem cell maintenance^57^, is reduced in *APP-KO* NPCs (Fig. 7a), with upregulation of canonical WNT inhibitors *DKK1, DKK3, KREMEN1, APCDD1* and *ALPK2* (Fig. 7b). We asked whether this reduction in WNT signaling might contribute to reduced NPC amplification in *APP-KO* NPCs. We compared control and *APP-KO* NPCs treated with the canonical ligand WNT3a for 48 hours to untreated NPCs and stained them for SOX2 and DCX (Fig. 7c-n). We observed a significant rescue of the loss of SOX2+ but no reduction in DCX+ cells (Fig. 7o,p), suggesting that APP regulates the balance between NPC proliferation and differentiation through partially parallel mechanisms, facilitating WNT signaling to promote NPC proliferation and repressing AP1 levels to prevent premature NPC differentiation.

**Fig. 7.**
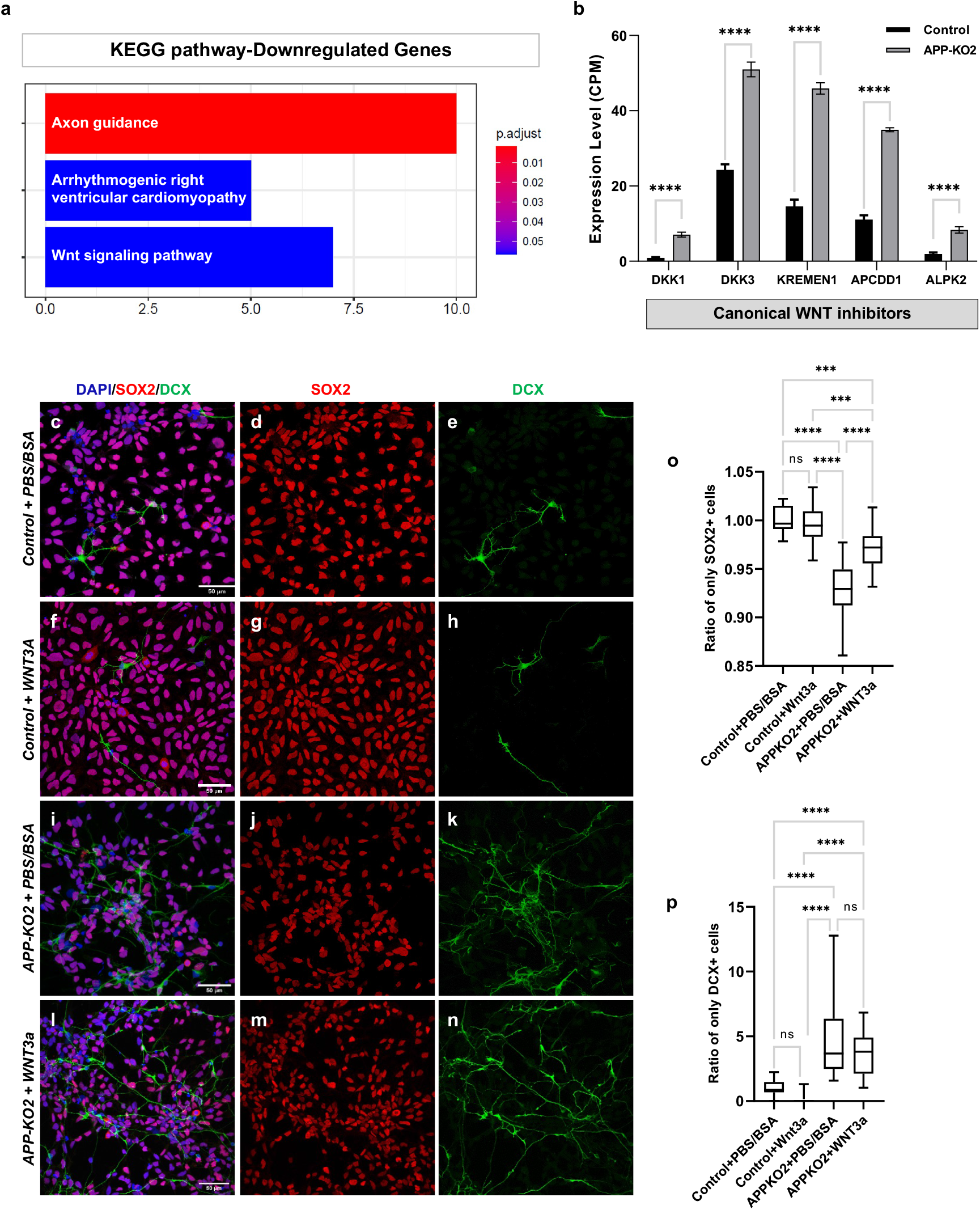
Loss of APP decreases amplification of NPCs by attenuating the canonical WNT pathway. **a**, KEGG pathway analysis shows downregulation of WNT signaling pathway in *APP-KO* background. **b**, Upregulation of canonical WNT inhibitors *DKK1, DKK3*, KREMEN1, *APCDD1*, and *ALPK2* in *APP-KO2* NPC compared to isogenic control (Ordinary one-way ANOVA, p<0.0001). **c-n**, Isogenic control and APP-KO2 NPCs treated with canonical WNT ligand (WNT3a) and stained for SOX2 and DCX. **o**, Ratio of SOX2+ was significantly rescued in *APP-KO2* NPCs treated with WNT3a (n=3, Ordinary one-way ANOVA, p<0.0001 for isogenic control+PBS-BSA vs *APP-KO2*+PBS-BSA and *APP-KO2*+PBS-BSA vs *APP-KO2*+WNT3a, p=0.0001 for isogenic control+PBS-BSA vs *APP-KO2*+WNT3a, p=0.0006 for isogenic control+WNT3a vs *APP-KO2*+WNT3a). **p**, Ratio of DCX+ cells was not rescued after treatment with WNT3a (n=3, Ordinary one-way ANOVA, p<0.0001 for all the significant conditions).

## Discussion

Here, we report what is, to our knowledge, the first evidence for a genetic network that specifically regulates the absolute timing at which human cortical progenitors generate neurons. Our evidence suggests that this timing requires simultaneously sufficient levels of canonical WNT signaling to promote progenitor identity as well as sufficient repression of AP1 activation to inhibit neurogenesis. We find that these two processes are genetically coupled by the activity of the Amyloid Precursor Protein. Our evidence supports a model in which this genetic network is required for the prolonged maintenance of human cortical NPCs in a progenitor state. In this model, loss of APP reduces WNT signaling and activates AP1 thus lowering the threshold for NPCs to transition from a self-renewing progenitor state to a neurogenic state. Strikingly, NPCs lacking APP generate cortical neurons in the correct temporal order. This means that the absolute amount of time a human cortical progenitor takes to generate neurons can be genetically uncoupled from the relative temporal order of the generation of cortical neurons. Obviously, evolution has achieved such uncoupling in the timing of cortical development in different mammalian species, where the same subtypes of neurons are generated in the same relative temporal sequence despite massive differences in the total amount of time required for cortical neurogenesis. Our findings suggest either that loss of APP mimics this evolutionary uncoupling, or, more excitingly, that regulation of genes in the APP-WNT-AP1 network may be part of the evolutionary uncoupling mechanism. In addition, whether the greater requirement for APP in humans compared to rodents may offer a clue into fAD is worth further investigation. We noted upregulation of the *MAPT* gene (1.36 Log2FC, Supplementary Table 3), encoding the other key AD protein Tau, in *APP* mutant progenitors. It is tempting to speculate that fAD mutations in *APP* have subtle effects on early human cortical development that cause reduced cortical robustness and resistance to a lifetime of neuronal and glial stress partially contributing to premature neurodegeneration.

Three observations suggest that human brain development may be particularly sensitive to loss of APP. First, there is a strongly reduced tolerance to complete loss of APP in human genetic data^29,58– 60^. Second, both *in vivo* and *in vitro* studies on mouse NPCs lacking APP have shown no consistent evidence for altered neurogenesis^8,35,36^. Third, among the genes whose expression is altered in *APP-KO* NPCs, 33 have been reported to be located within human accelerated regions (HARs), which are conserved genomic loci with elevated divergence in humans^61,62^. We also did not observe any effect on human motor neurogenesis, suggesting that APP function may be more important during cortex development. Because APP is highly conserved in sequence and expression among mammalian species, we speculate that the striking requirement for APP in human cortical neurogenesis is less due to anything special about the human APP protein itself, and more due to the special features of human cortical neurogenesis. In particular, we propose that the links between APP, WNT and JNK pathways have been co-opted in human, and perhaps more generally primates, to prolong the ability of the cortical NPCs to retain a progenitor state while also generating more differentiated daughters. For example, human/primate cortical NPCs may express human/primate-specific regulators of APP activity or stability that allow them to suppress JNK signaling and enhance WNT signaling more efficiently than in other mammalian species. Alternatively, human cortical NPCs may be particularly sensitive to cellular stress. Some studies suggest that increased stress drives premature neurogenesis in cortical progenitors^63^. APP is a conserved stress response protein, as are JUN and FOS. Therefore, the absence of APP may mimic cellular stress, activate AP1-mediated responses, and drive premature neurogenesis. We have previously shown that the coding sequence contributes to evolutionary changes in the neurogenic activity of proneural proteins for example^64^. We cannot exclude that subtle changes to the otherwise highly conserved APP coding sequence may be involved in its acquisition of this function in human cortical NPCs.

Increasing evidence points to human specific genetic alterations, such as new gene isoforms ^33,65– 68^, microRNAs ^69^ and regulatory sequences, creating human-specific transcriptional networks ^61,70^ potentially resulting in human specific brain features. A tantalizing speculation is that members of the genetic network we uncovered here may be regulated by one or more human specific protein isoform or microRNA whose expression is enriched in the cortex. Interestingly, a clinical case of a homozygous nonsense truncating mutation in APP leading to microcephaly has been reported^59,60^. The reported patient carries mutations in three genes including homozygous mutations in *APP*, that strongly reduce its expression levels, and *SETX* as well as a heterozygous mutation in *CLN8*. Since *CLN8* is associated with autosomal-recessive neuronal ceroid lipofuscinosis (MIM: 600143) and the patient is heterozygote for that, the authors conclude that the phenotype is not related to this gene. *SETX* is associated with autosomal-dominant juvenile amyotrophic lateral sclerosis type 4 (ALS; MIM: 602433) and autosomal recessive ataxia with ocular apraxia type 2 (AOA2; MIM: 606002). Since ALS and AOA2 usually have onset in adolescence, they conclude that it is more probable that the phenotype is related to the mutation of *APP* and not *SETX*. This supports the notion that human brain development is vulnerable to loss of APP and that perhaps the status of *APP* heterozygosity should be included in clinical genetic counseling.

**Extended Data Fig. 1.**
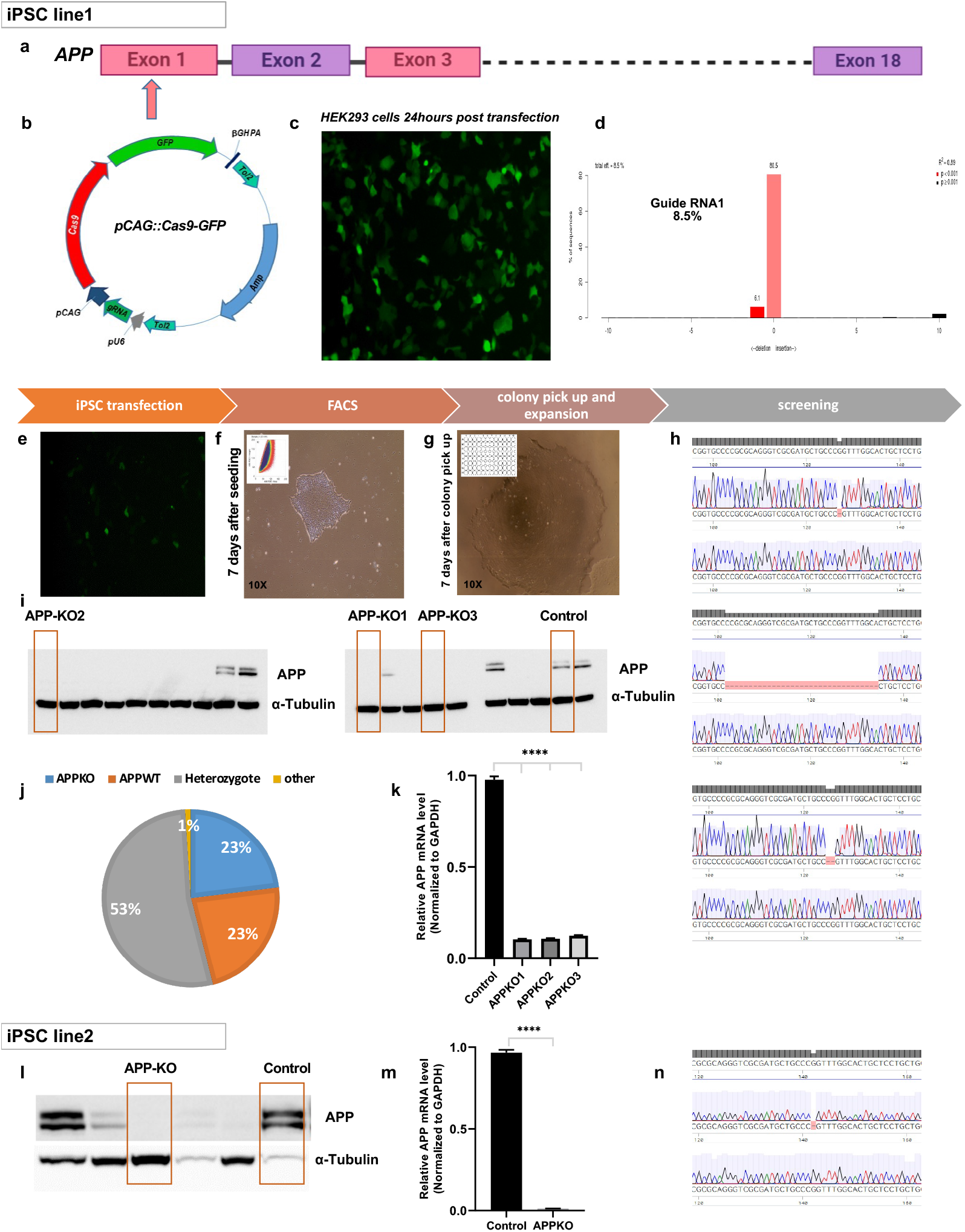
Generation of iPSC derived APP knockout clones. **a-k**, Individual steps in generating *APP* knock out from iPSC line WTSIi002-A, henceforth “Line 1”. **a**, Targeting exon1 of the *APP* gene by **b**, a plasmid vector containing the guide RNA, CAS9 and GFP. **c**, HEK293 cells 24 hours after transfection. **d**, Cleavage efficiency of Guide RNA1 that was used for transfection of iPSC line 1. **e-h**, iPSC transfection by Guide RNA1, post FACS morphology and sequencing results of clones. **i**, Western blot results showing undetectable APP protein in 14 out of 60 clones. One isogenic control transfected with Cas9 and the guide RNAi but not mutated for APP and three *APP-KO* clones were chosen for further experiments. **j**, Distribution of clones with different *APP* genotypes (23%=*APP-KO*, 23%=*APPWT* and 53%=heterozygote and 1%=other mutation). **k**, qPCR confirms low level of *APP* mRNA expression in *APP-KO* clones. **l-n**, Generation of *APP-KO* clones from iPSC line WTSIi008-A, henceforth “Line 2”, confirming very low level of APP mRNA and undetectable APP protein.

**Extended Data Fig. 2.**
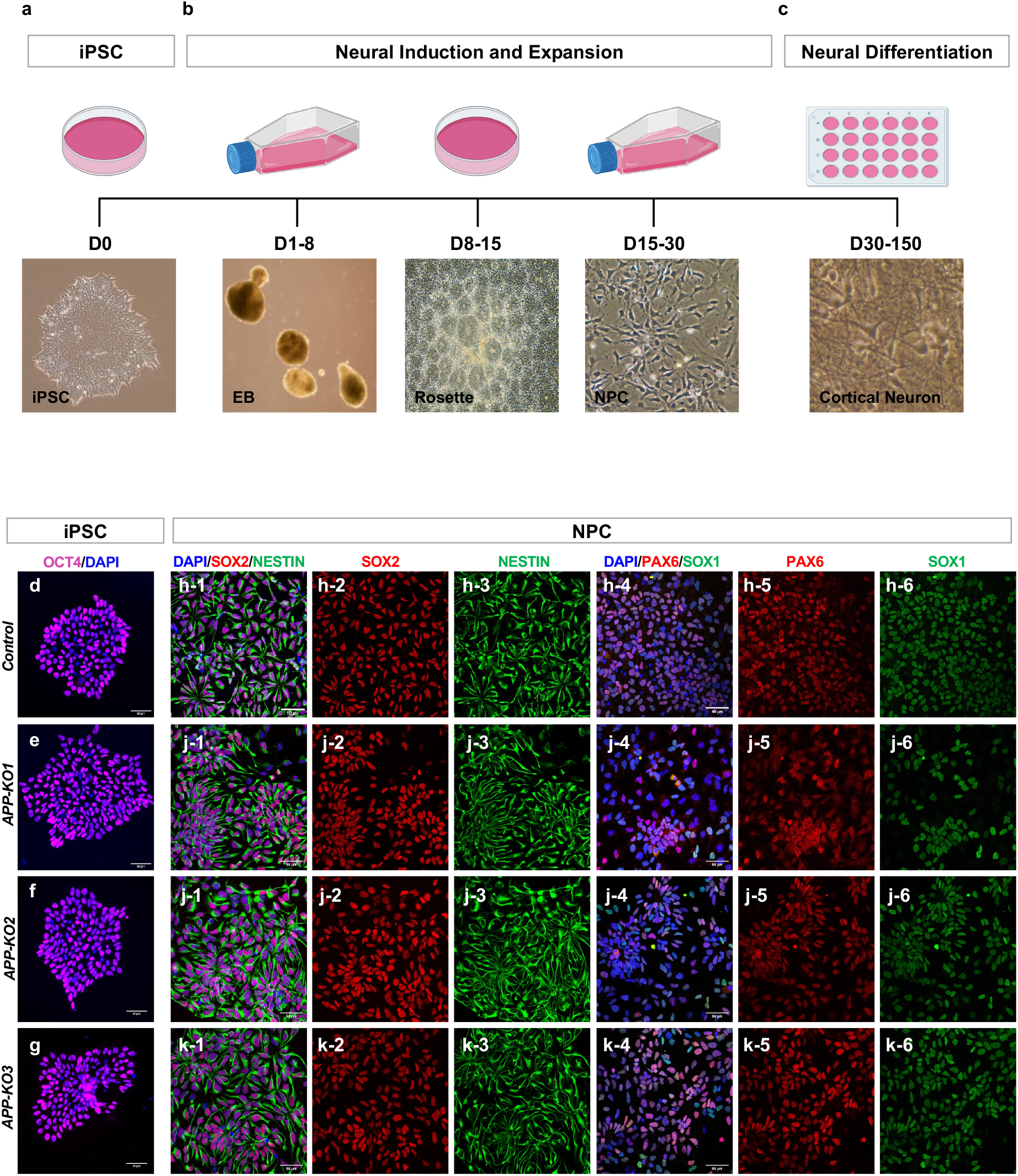
Timeline of the cortical differentiation protocol and characterization of neural progenitor cells. **a-c**, Schematic showing different steps in generating cortical neurons (culture tools adapted from icons by BioRender.com). **d-g**, Confirming pluripotency of isogenic control and *APP-KO* iPSCs with pluripotency marker OCT4. **h1-k6**, Characterization of isogenic control and *APP-KO* derived neural progenitor cells with markers: Nestin, SOX2, PAX6, SOX1 (scale bar 50µm).

**Extended Data Fig. 3.**
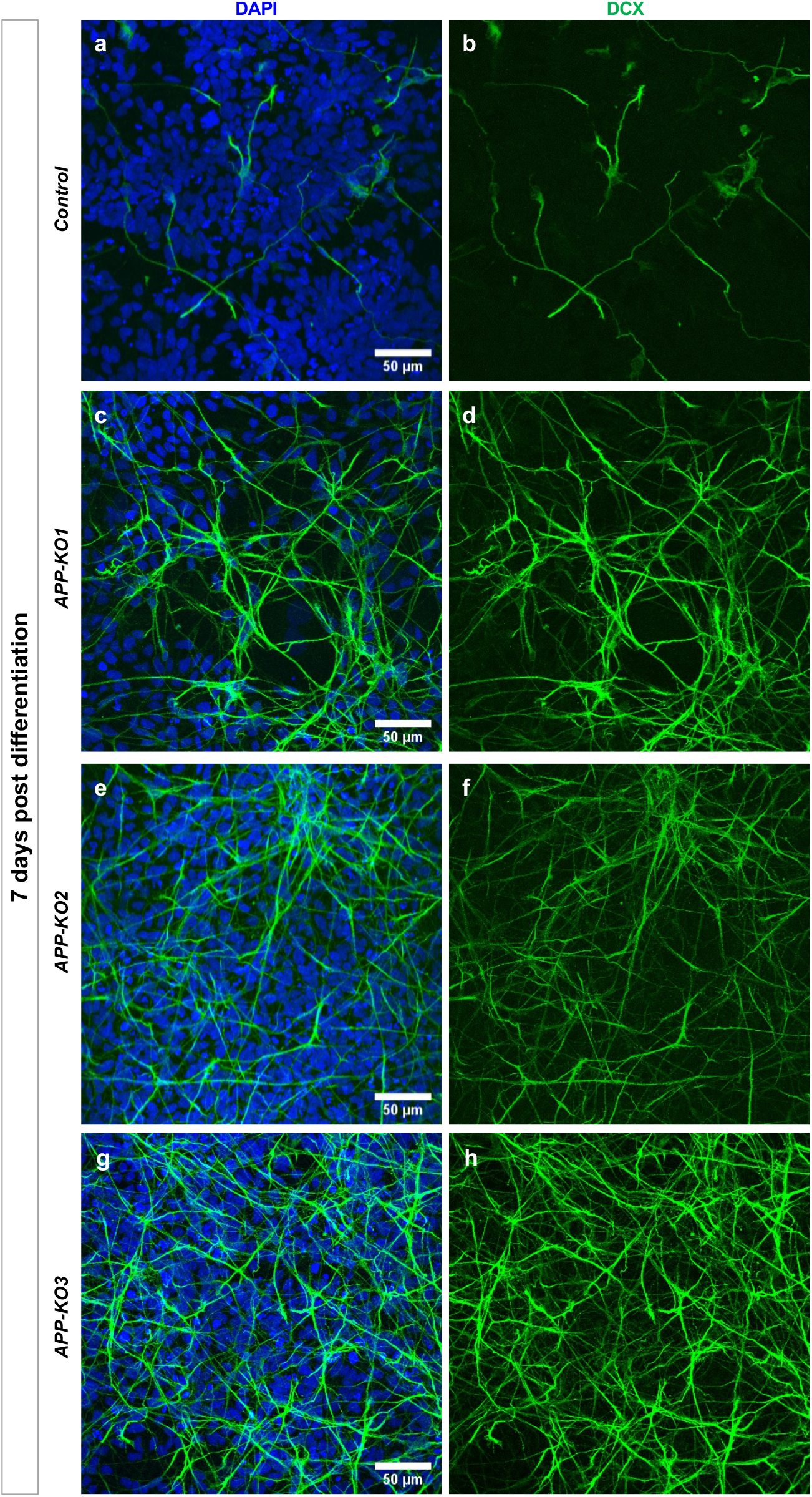
Increased production of neuronal precursors and young neurons in *APP-KO* NPCs 7 days after onset of neuronal differentiation. **a-h**, An increase in the number of cells expressing the newly born neuron marker Doublecortin (DCX) in *APP-KO* background compared to isogenic control 7 days post-differentiation (Scale bar 50 µm).

**Extended Data Fig. 4.**
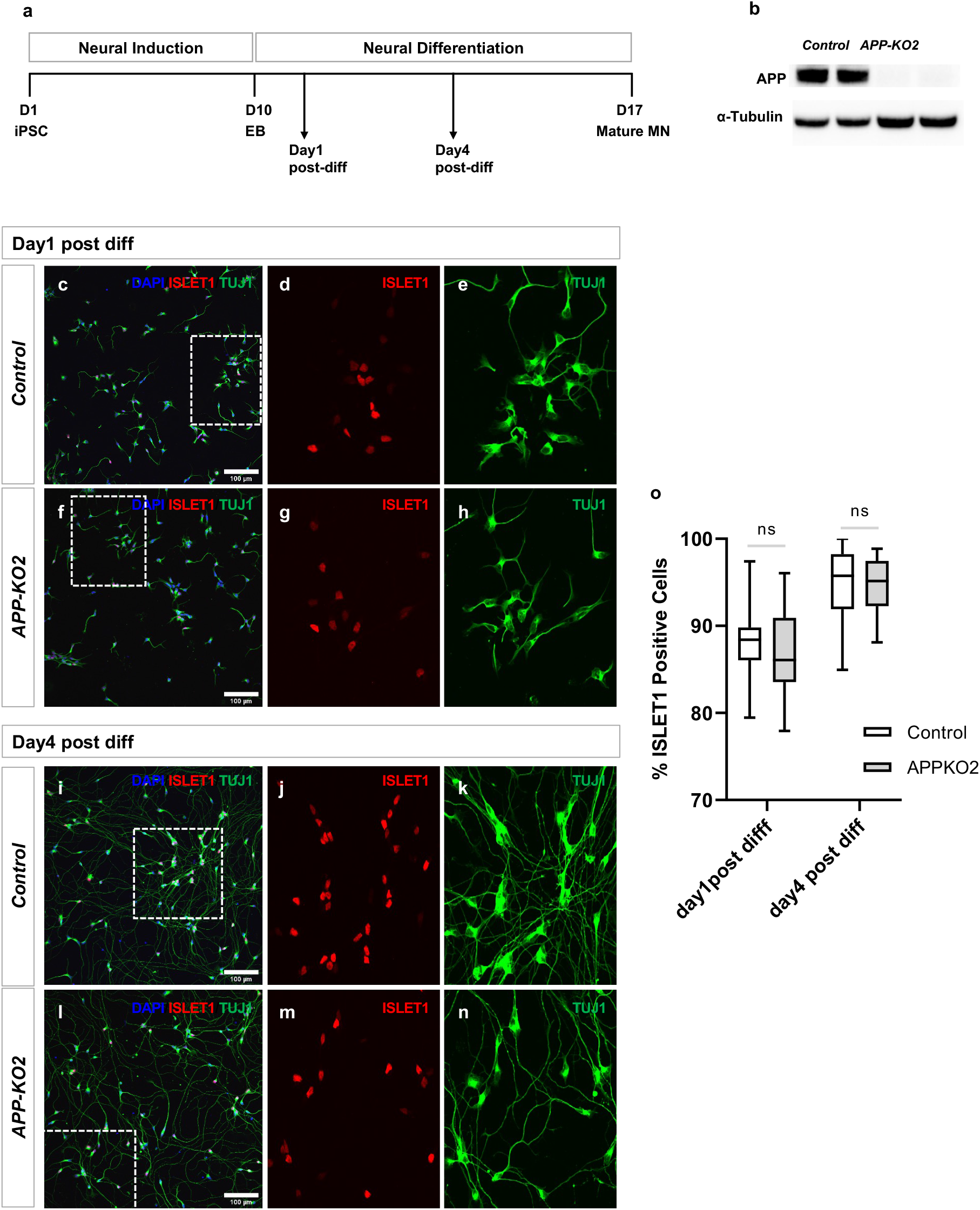
No difference in the number of ISLET1+ motor neurons derived from isogenic control and *APP-KO2*. **a**, Schematic showing the timeline of different steps in generating motor neurons. **b**, APP expression was confirmed in isogenic control motor neuron progenitors by western blot. **c-h**, Day1 and **i-n**, Day4 post-differentiation motor neurons stained for ISLET1 and TUJ1 (scale bar 100µm). **o**, No significant differences were observed in the percentage of ISLET1+ cells at 2 different time points, day1 and day4 post-differentiation (n=3 biological independent repeat, 2way ANOVA).

**Extended Data Fig. 5.**
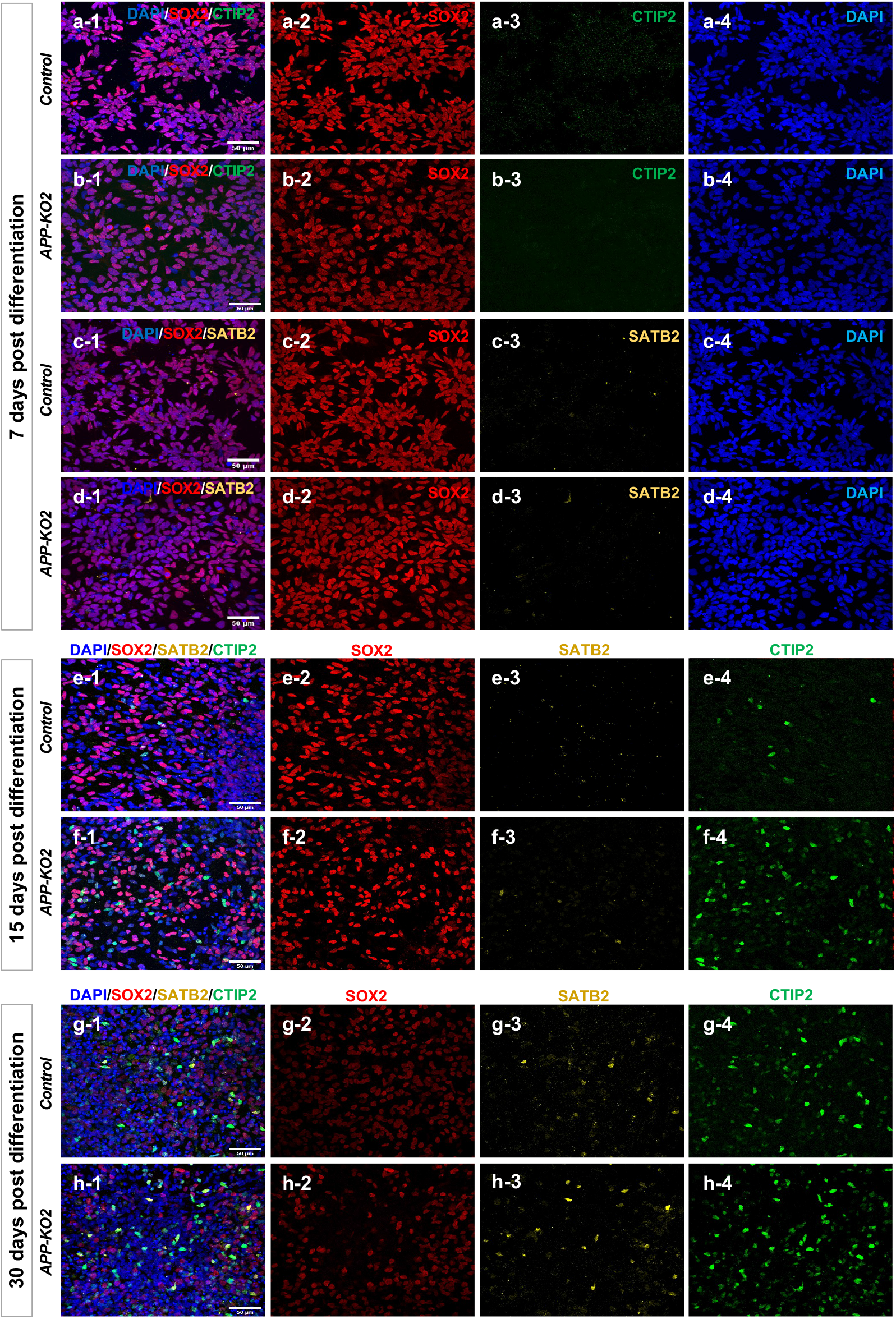
Cortical neuron fate markers appear with same temporal order in *APP-KO* and isogenic control. **a1-d4**, No expression of CTIP2 (early born neuron) and SATB2 (late born neuron) was observed 7 days post-differentiation. **e1-f4**, CTIP2 and SATB2 appeared at the same time 15 days post-differentiation in isogenic control and *APP-KO2*. **g1-h4**, Expression of CTIP2 and SATB2 30 days post-differentiation in isogenic control and *APP-KO2* (scale bar 50µm for all images).

**Extended Data Fig. 6.**
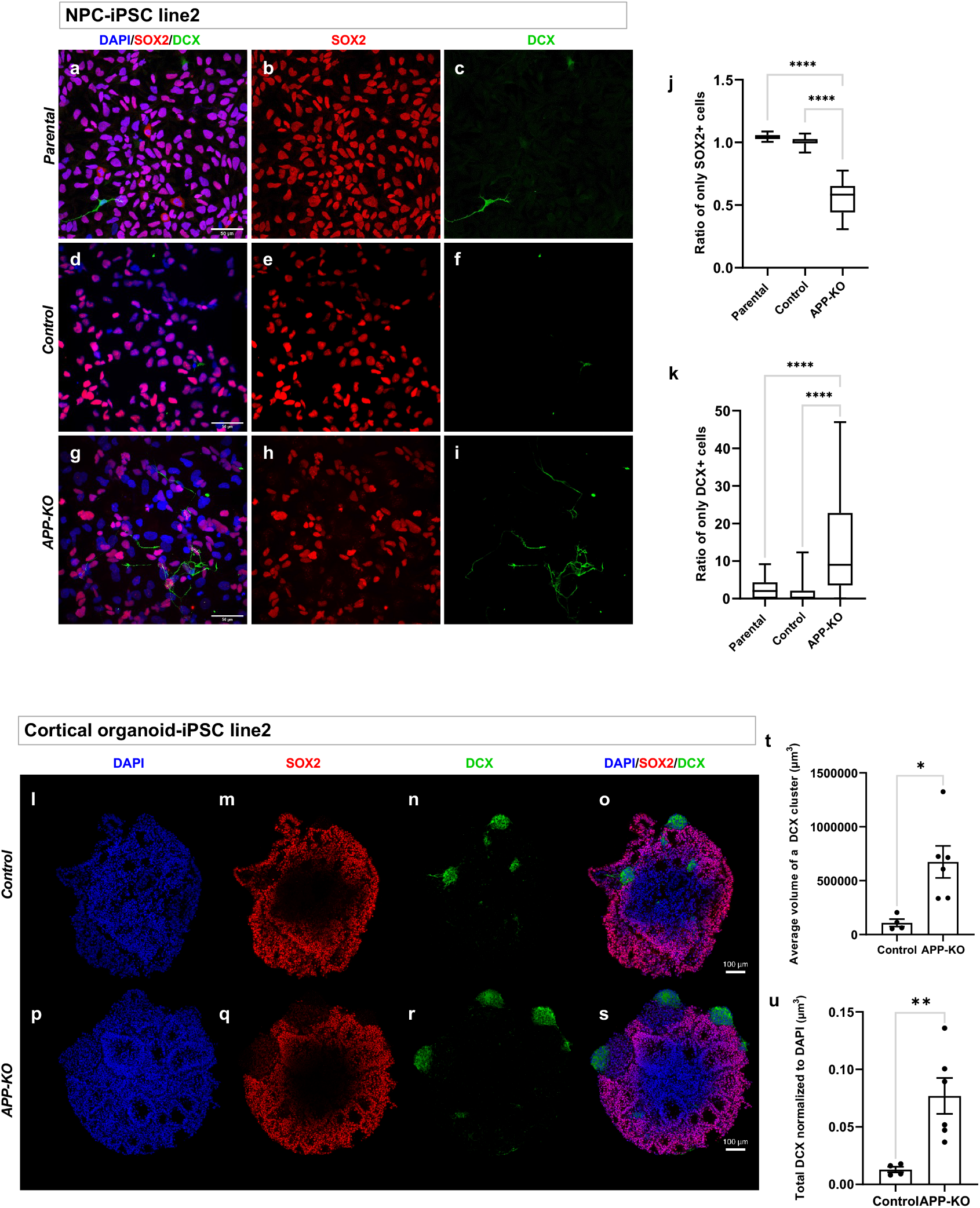
Premature differentiation due to loss of APP is reproducible in NPC and cortical organoid from a different genetic background. **a-i**, NPCs from parental, isogenic control (i.e. transfected with guide RNA and Cas9 but not mutated for APP) and APP-KO from iPSC line 2 stained for SOX2/DCX. **j**, Quantification of SOX2+ cells and **k**, DCX+ in NPCs showed that the results were reproducible independent from genetic background (n=3, Ordinary one-way ANOVA, p<0.0001, scale bar 50µm). **l-s**, Cortical organoid for isogenic control and *APP-KO* from iPSC line 2 stained for SOX2/DCX at day 15 of culture showing **t-u**, significant increase in total DCX (p=0.0089) and average volume of a DCX cluster (p=0.0115) in APP-KO background compare to isogenic control (n=4 organoids for isogenic control and n=6 organoids for APP-KO, scale bar 100µm, supplementary movies I and II).

**Extended Data Fig. 7.**
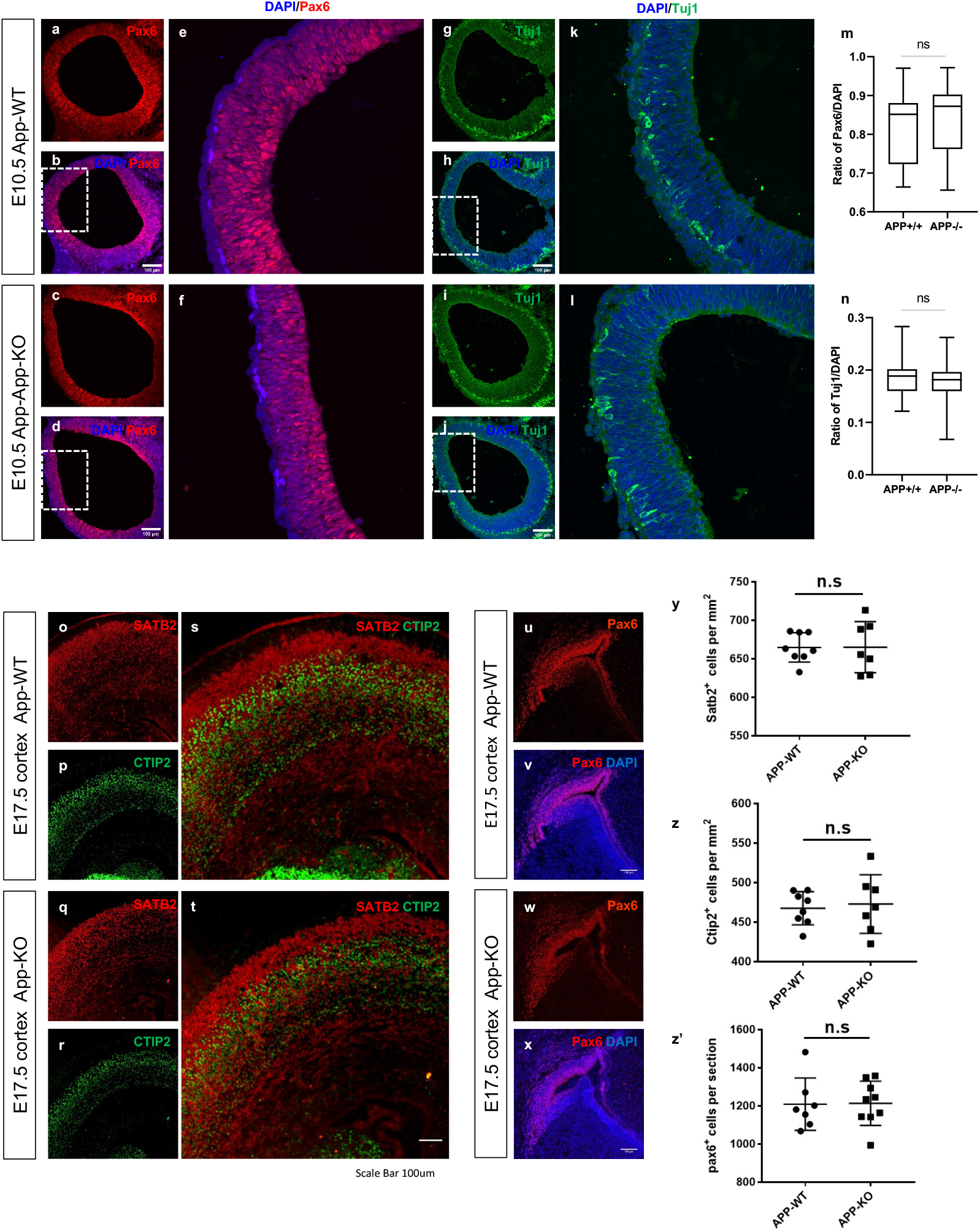
Normal cell type numbers during mouse cortical neurogenesis in the absence of APP. **a-l**, Brain sections of E10.5 embryo stained for Pax6 and Tuj1. **m-n**, No difference was observed in the ratio of Pax6/DAPI and Tuj1/DAPI in *APP-WT* and *APP-KO* embryo (n=3 embryos, unpaired t-test, scale bar 100µm). **o-x**, Brain sections of E17.5 embryo stained for Ctip2, Satb2 and Pax6. **y-z’**, No difference was observed in the number of Satb2+ and Ctip2+ cells (n=8 for *APP-WT* and n=7 *APP-KO*, unpaired t-test, scale bar 100µm) and number of Pax6+ (n=7 for *APP-WT* and n=9 *APP-KO*, unpaired t-test, scale bar 100µm).

**Extended Data Fig. 8.**
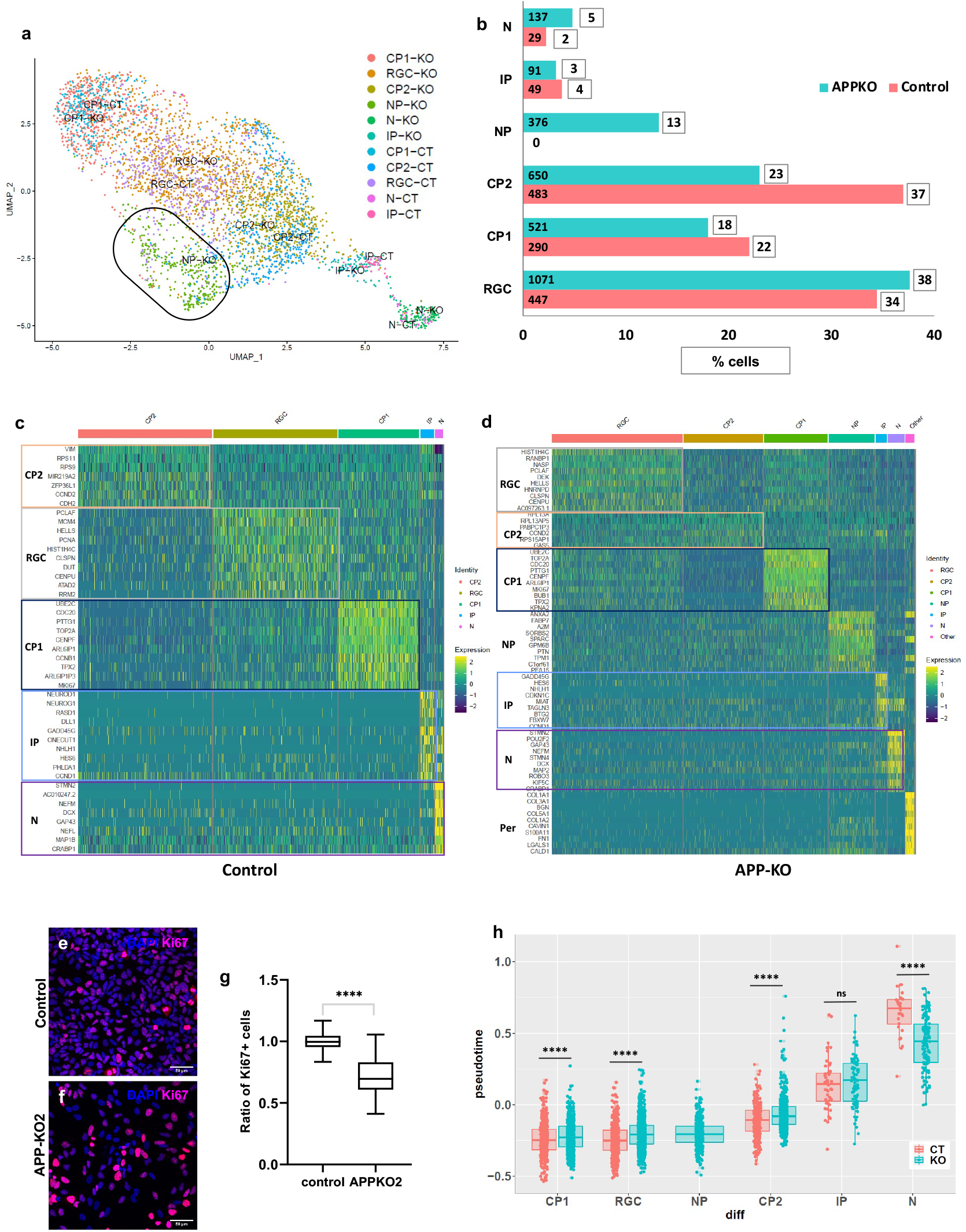
APP-KO NPCs are in a temporally advanced neurogenic state. **a**, UMAP of clustering according to the cell identity shows different clusters (RGC=radial glial cell, CP1=cycling progenitor1, CP2=cycling progenitor2, IP=intermediate progenitor, NP=neurogenic progenitor, N=neuron). **b**, Number and percentage of cells in each cluster in isogenic control vs *APP-KO2* **c-d**, Heat maps of different clusters in isogenic control and *APP-KO2*. **e-g**, Significant decrease in ratio of Ki67+ cells in *APP-KO2* (n=3, unpaired t-test, p<0.0001, scale bar 50µm). **h**, Pseudotime analysis shows a shift toward neuronal fate in *APP-KO* clusters (p-value = 5.235e-10 for RGC, p-value = 4.554e-05 for CP2, p-value = 0.01949 for CP1, p-value = 0.1675 for IP, p-value = 1.457e-06 for neuron, Two-sample Kolmogorov-Smirnov test is used for comparison of all the clusters).

**Extended Data Fig. 9.**
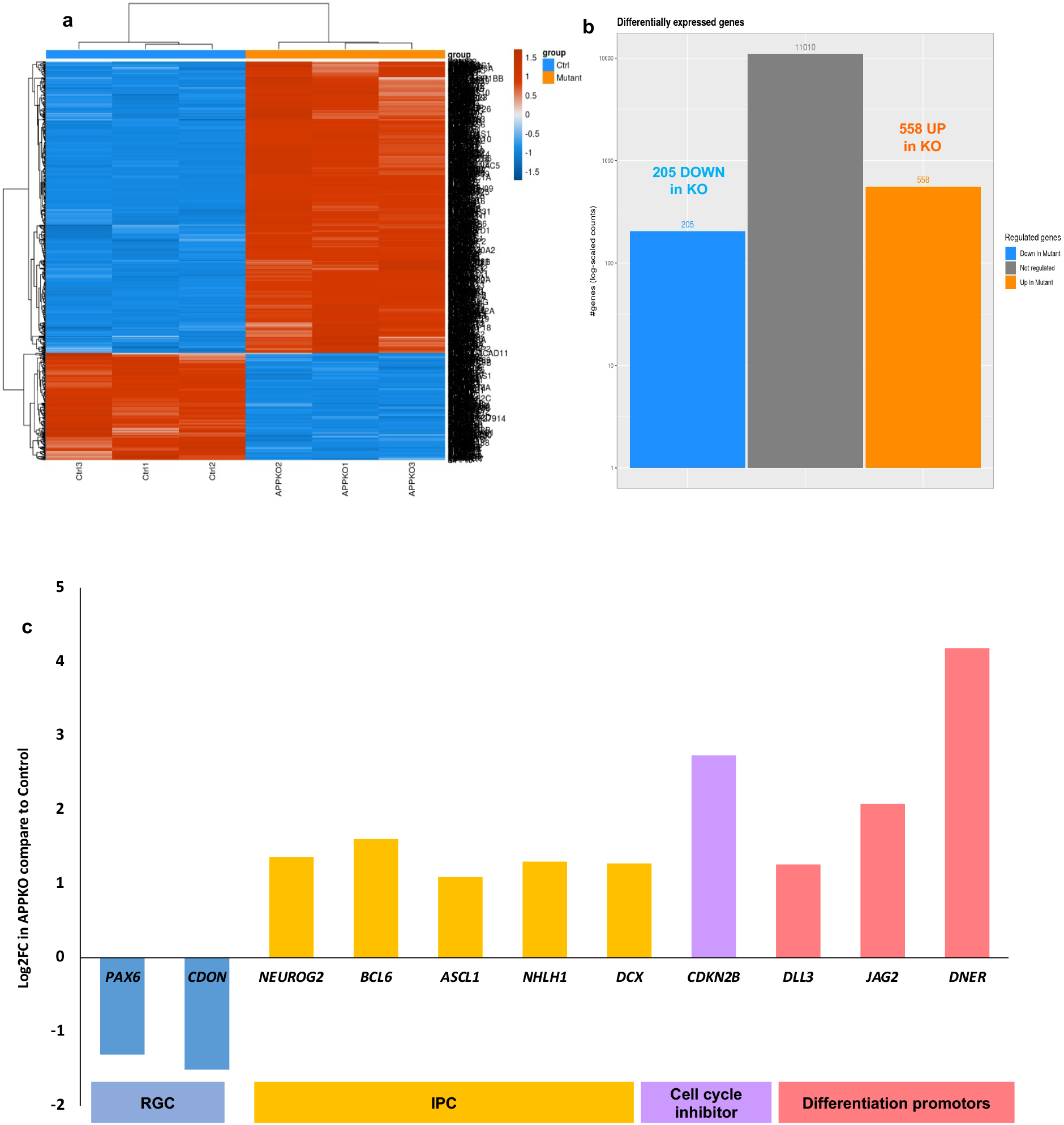
Bulk RNA seq confirms the shift towards a differentiated fate in APP-KO NPCs. **a**, Heat map of differentially expressed genes shows strikingly different patterns in isogenic control and *APP-KO2* NPCs. (*Ctrl1*,2,3 correspond to 3 technical repeats of isogenic control and *APP-KO1,2,3* correspond to 3 technical repeats of *APP-KO2*). **b**, 763 differentially expressed genes in *APP-KO2* NPCs compared to isogenic control which 558 genes are upregulated and 205 gens are downregulated. **c**, Bulk RNAseq shows downregulation of RGC markers (PAX6 and CDON) and upregulation of neurogenic genes (*NEUROG2, BCL6, ASCL1, NHLH1*, and *DCX*), cell cycle inhibitor (*CDKN2B*) and differentiation promoting Notch ligands *DLL3, JAG2, DNER*.

## Material and Methods

### iPSC culture and maintenance

iPSC cell lines WTSIi002-A (https://cells.ebisc.org/WTSIi002-A/, iPSC line1) and WTSIi008-A (https://cells.ebisc.org/WTSIi008-A/, iPSC line2) were purchased from EBISC (European Bank for Induced pluripotent Stem Cells) and maintained on Geltrex LDEV-Free hESC-qualified Reduced Growth Factor Basement Membrane Matrix (ThermoFisher Scientific) in Essential 8™ Flex Media Kit (ThermoFisher Scientific) with 0.1% penicillin/streptomycin. Cultures were fed every other day and passaged every 5–7 days by ReLeSR™ (STEMCELL Technologies).

### Creating *APP-KO* clones by CRISPR/Cas9

#### iPSC line1

Four guide RNAs were designed by Crispor (http://crispor.tefor.net/) to target APP’s first exon and cloned in pCAG-CAS9-GFP (Extended Data Fig. 1, A-B). To evaluate the cleavage efficiencies of the guide RNAs, HEK293 cells were transfected with p-CAG-Cas9-GFP containing four guide RNAs (Extended Data Fig. 1C). DNA extraction and PCR were performed 48 hours post transfection for the cleaved region. Sequenced PCR products were analyzed in TIDE (Tracking of Indels by Decomposition) to obtain cleavage efficiency. The sequence of each guide RNA, their cleavage efficiency, and off-target are listed in Supplementary Table 5. The best guide RNA with 8.5% total efficiency and lowest off-target was chosen for further experiments (Extended Data Fig. 1D). Karyotyping was performed on the iPSC line (which we refer to as “parental line” 1) before the CRISPR approach and the results showed a deletion in chromosome 18. The iPSC line was transfected with the best guide RNA using Lipofectamine stem reagent (ThermoFisher Scientific). GFP-positive cells were isolated by FACS 48 hours post-transfection and seeded at low density. A total of 118 clones were picked after 5 days, expanded, and 60 screened for possible mutations (Extended Data Fig. 1E-1H). 16 out of the 60 clones had homozygous mutations for APP. These mutations are classified as: 1, 2, 3, 20, 22, 33bp deletions and 1bp and 2bp insertions (Supplementary Table 6). The absence of expression of the APP protein was confirmed by western blot in 14 clones (Extended Data Fig. 1I). The efficiency of producing APP knock-out clones was 23% (Extended Data Fig. 1J). Three APP knock-out clones were chosen for further experiments and low APP mRNA levels were confirmed by quantitative-PCR (Extended Data Fig. 1K). The control, which was chosen for further experiments, expressed guide RNA and CAS9 but the APP gene remained un-mutated. Karyotyping was performed for the APP-KO clones, control, and parental iPSC. The results showed that the same deletion in chromosome 18 in parental line was also confirmed in control and *APP-KO* clones. No other deletion or duplication was observed in *APP-KO* clones and control after CRISPR approach. The pluripotency of *APP-KO* cells and the control were confirmed by OCT4 staining (Extended Data Fig. 2D-2G).

#### iPSC line2

Karyotyping was performed on the iPSC line before the CRISPR approach (which we refer to as “parental line” 2) and the results showed a duplication in chromosome 20. The guide RNA 1 and the ribonucleoprotein approach were used to generate *APP-KO*. 1×10^6^ iPSCs were nucleofected with the RNP complex from IDT (225pmol of each RNA and 120pmol of Cas9 protein). iPSCs were plated 48 hrs later at very low density (10 cells/cm^2^) on Laminin-521 with CloneR supplement (Stem Cell Technology) for clonal selection. One week later, iPSC clones were picked under a stereomicroscope and cultured on Laminin-521 in 96 well plates (Duscher). The resulting iPSC clones were duplicated after confluency and used for cryoconservation and DNA extraction. One base-pair deletion was observed in screening results and the absence of expression of APP protein was confirmed by western blot. The APP knock-out clone and control were used for further experiments and the low level of APP mRNA was confirmed by quantitative-PCR (Extended Data Fig. 1 L-N). Karyotyping was also performed for the *APP-KO* clone, control, and parental iPSC. The results confirmed the same duplication in chromosome 22 in all the conditions. No other deletion or duplication was observed in *APP-KO* clones and control after CRISPR approach.

### Cortical differentiation of human iPSC

The cortical differentiation protocol is divided into 2 main steps: neural induction and neural differentiation. At day 0, iPSCs were detached by Accutase (ThermoFisher Scientific) and transferred to T75 ultra-low attachment flasks (VWR) in Essential 8 medium with 0.1% penicillin/streptomycin and Stemgent hES Cell Cloning & Recovery Supplement (Ozyme, 01-0014-500) to form Embryoid body (EB). At day 1, the medium was switched to EB medium containing DMEM/F12 Glutamax, 20% knockout serum replacement, 1% nonessential amino acids, 1% penicillin/streptomycin, 0.55mM 2-mercaptoethanol, and supplemented with Dorsomorphin (1µM, Sigma-Aldrich P5499), SB431542 (10µM, Abcam ab120163) for 8 days. At day 8, Embryoid bodies (EBs) were collected and seeded on Geltrex and maintained for 6-8 days in Neurobasal without vitamin A medium and B27 without vitamin A supplemented with Human EGF (10ng/ml) and human FGF2 (10ng/ml). Cells were fed every day from day 0 to day 16. Rosettes were manually picked and dissociated with Accutase, seeded on poly-ornithine and laminin-coated dishes for expansion, and maintained with passage for two additional weeks to achieve a large pool of neural progenitor cells (NPCs) (Extended Data Fig. 2 A-C). Characterization for NPCs was performed by staining for Nestin, SOX2, SOX1, and PAX6 (Extended Data Fig. 2 H1-K6). NPCs were seeded on 24-well plates with coverslips and the day after seeding, the medium was switched to the B-27 Plus Neuronal Culture System (ThermoFisher Scientific) supplemented with Ascorbic acid. Cortical neurons were stained for early-born neural markers and late-born neural markers at the indicated early time points or kept for 3 to 4 months in culture and stained for synaptic markers at 60 days and 120 days post differentiation.

### Motor neuron differentiation and staining

Control and mutant iPSC clones were differentiated into motor neurons as described^31^. In summary, after dissociation of EBs containing motor neuron progenitors at D10, single cells were plated on poly ornithine-laminin coated coverslips at 2×10e5 cells/coverslip for 1, 4, and 7 more days. Cells were fixed with 4% paraformaldehyde in PBS for 10 minutes RT. For staining, cells were treated with 5 % goat serum with 0.1% Triton X100 (Thermofisher Scientific) for 1hour at RT and then incubated overnight at 4°C with a mouse anti-βIII-Tubulin (TUJ1, Sigma, 1/500) and a rabbit anti-ISLET1 (Abcam, 1/100) antibodies in 5% goat serum. Secondary antibodies (goat anti-mouse IgG2a-Alexa 555 (1/2000) and goat anti-rabbit IgG Alexa 488 (1/2000) from ThermoFischer Scientific) were incubated for one hour at RT with Hoechst33342 to stain nuclei.

### Sparse labeling and racing of NPCs

Sparse labeling was achieved using a lentiviral viral vector approach as described^32,33^. *APP-KO* and control NPCs were seeded on Poly-L-Ornithine/Laminin coated coverslips and transduced by virus Lenti-Synapsin-GFP. The medium was changed to fresh Neurobasal without vitamin A and B27 without vitamin A supplemented with EGF and hFGF2 to wash out lentiviruses one day after initial infection. When the GFP+ cells appeared, it was considered as day 0 and the medium was switched to differentiation medium (B-27 Plus Neuronal Culture System) and kept for 7 and 30 days. Lentiviral expressing cells were detected by anti-GFP antibody and cell type was determined by anti-SOX2 and anti-TUJ1 antibodies.

### Immunocytochemistry

Cells were fixed in PFA 4% and incubated for 30 min in blocking buffer (phosphate-buffered saline and Triton X-100 0.3% and 5% horse serum). Primary antibodies (Supplementary Table 7) were diluted in antibody solution (phosphate-buffered saline and Triton X-100 0.3% and 5% horse serum) and applied overnight at 4°C. After three washes in phosphate-buffered saline, secondary antibodies conjugated to Alexa fluorophores (Molecular Probes, Eugene, OR, USA) were diluted at 1:1000 in blocking buffer and applied for 1 hr at room temperature. Cells were washed in phosphate-buffered saline followed by nuclear staining by DAPI (Sigma) for 10 min. Cells were washed three more times and mounted by VECTASHIELD® Mounting Medium-Vector Laboratories. Confocal image acquisition was performed using Olympus FV1200 and SP8 Leica DLS.

### Generation of cortical organoids and immunohistochemistry

Cortical organoids were generated from human iPSCs using a previously reported protocol^71^, with some modifications. hiPSCs were incubated with Accutase (Life Technologies, A1110501) at 37 °C for 7 min and dissociated into single cells. In order to obtain uniformly sized spheroids, approximately 3 × 10^6^ single cells were added per well in the AggreWell 800 plate (STEMCELL Technologies, 34815) with Essential 8 flex medium supplemented with Stemgent hES Cell Cloning & Recovery Supplement (1X, Ozyme, STE01-0014-500) and incubated at 37°C with 5% CO_2_. After 24h, spheroids from each microwell were collected by firmly pipetting medium in the well up and down and transferred into Corning® non-treated culture dishes (Merck, CLS430591-500EA) in TeSR™-E6 (StemCell Technologies, #05946) supplemented with two inhibitors of the SMAD signaling pathway, dorsomorphin (2.5 μM, Sigma-Aldrich, P5499) and SB-431542 (10 μM, Abcam, ab120163). From day 2 to day 5, TeSR™-E6 supplemented with dorsomorphin and SB-431542 was changed every day. On the sixth day in suspension, neural spheroids were transferred to neural medium containing Neurobasal minus vitamin A (Life Technologies, 10888), B-27 supplement without vitamin A (Life Technologies, 12587), GlutaMax (1%, Life Technologies, 35050), Penicillin-Streptomycin (1%, Life technologies, 15140122) and 2-mercaptoethanol (5mM, Life technologies, 31350010). The neural medium was supplemented with 10 ng/mL epidermal growth factor (PreproTech, AF-100-15) and 10 ng/mL basic fibroblast growth factor (R&D Systems, 234-FSE-025) and changed daily until day 12. From day 12 to day 24, medium was changed every other day. At day 15, spheroids were fixed with 2% paraformaldehyde at 4°C for 6h and rinsed 3 times with PBS for 10 minutes. Following 2 washing with PBS + 2% Triton-X100 for 2 hours, spheroids were treated overnight at RT with PBS + 2% Triton-X100 + 2% Tween20 + 20%DMSO. Blocking/permeabilization was performed with PBS + 10% Horse serum, 3% BSA, and 2% Triton-X100 for 24h at RT. Primary antibodies, (Sox2 1/500, Millipore AB5603) and (DCX 1/2000, Millipore AB2253), were incubated in the same solution supplemented with 0.05% Azide for at least 3 days at 4°C. After multiple washing with PBS + 0.5% Tween-20 until the next day, organoids were then incubated for at least 3 days with secondary antibodies. After 2 days of washing with PBS + 0.5% Tween-20, samples were cleared overnight in RapiClear 1.49 (SunJin lab) before mounting on cavity slides (Hecht Karl™ 42410010).

### Doublecortin (DCX) quantification

Whole-mount day 15 organoids were imaged with a Nikon A1R HD25 confocal microscope at 1µm intervals with a 10X objective (MRD71120). All analyses on organoids were performed using the NIS-Elements software (Nikon). Image stacks were reconstructed in 3D to measure volume of DCX clusters and total DAPI. A threshold of 3000 µm^3^ was fixed to only quantify DCX clusters. Each measurement was normalized to the total volume of DAPI in the organoid. Each organoid is considered as n=1.

### Embryo collection, immunohistochemistry, and antibodies

E10.5 and E17.5 embryos were collected from *App-WT* or *App-KO* pregnant mice, whole embryos were fixed in 2% paraformaldehyde (PFA) in PBS at 4 °C for 2-3 hours, then dehydrated in 30% sucrose in 1X PBS overnight (o/n). After all the samples sank into the bottom of the tube, they were embedded in OCT compound (TissueTek) and frozen at -20 °C. Sagittal sections were performed by cryostat (Leica) at 20 μm and then slices stored at -80 °C. For the immunostaining, sections were fixed with 4% PFA for 10 minutes at RT, then blocked with 10% normal donkey or goat serum in 1 X PBS with 0.1% Triton (PBT) for 1 hour at RT followed by 3 washes in 1 X PBT. Thereafter, these sections were incubated with primary antibodies diluted in 0.1% 1 X PBT containing 1% normal donkey or goat serum o/n at 4 °C or 3-4 hours at RT. After 3 washes with 1X PBT, samples were incubated with appropriate secondary antibodies conjugated with Alexa Fluor 488, Alexa Fluor 555, or Alexa Fluor 647 (1:500, Invitrogen) in 0.1% 1 X PBT containing 1% normal donkey or goat serum for 1-2 hours at RT. After washing with 1X PBT for 3 times, then counterstained with DAPI (1:2000, Sigma), the slides were mounted by using Vectashield (Vector) after rinsing. After staining, images were obtained by using confocal microscope Olympus FV-1200.

### qPCR

The cells were lysed directly in the wells by addition of 300 μl Buffer RLT supplemented with 15 mM beta-mercaptoethanol (Thermo Fisher Scientific) after a wash with Dulbecco’s Phosphate-Buffered Saline (DPBS, Life Technologies). Total RNA was isolated using the RNeasy Mini extraction kit (Qiagen, Courtaboeuf, France) according to the manufacturer’s protocol. RNA levels and quality were quantified using a Nanodrop spectrophotometer. cDNA synthesis was performed by Thermo Scientific Verso cDNA Synthesis Kit and Quantitative PCR assay was performed by Sybergreen Gene Expression Assays in triplicate wells of 96-well plates. Primers are listed in Supplementary Table 8 and 2^-(Cp GOI – Cp internal gene)^ was used for analysis. The housekeeping gene (GAPDH) was selected to control for variation in cDNA amounts.

### Bulk RNA sequencing and analysis

The cells were lysed directly in the wells by addition of 300 μl Buffer RLT supplemented with 15 mM beta-mercaptoethanol (Thermo Fisher Scientific) after a wash with Dulbecco’s Phosphate-Buffered Saline (DPBS, Life Technologies). Total RNA was isolated using the RNeasy Mini extraction kit (Qiagen, Courtaboeuf, France) according to the manufacturer’s protocol. RNA levels and quality were quantified using a Nanodrop spectrophotometer. RNA sample purity/integrity was assessed using an Agilent 2200 Tapestation. mRNA library preparation was completed following the manufacturer’s recommendations (KAPA mRNA hyperprep ROCHE). Final samples pooled library prep were sequenced on Nextseq 500 ILLUMINA with MidOutPut cartridge (2×130Million 75 base reads) with 1 run, corresponding to 2×20Million reads per sample after demultiplexing. The quality of raw data was evaluated with FastQC. Poor quality sequences were trimmed or removed with fastp software to retain only good quality paired reads without adapters. Star v2.5.3a^72^ was used to align reads on the hg19 reference genome using standard options. Quantification of gene and isoform abundances was carried out with rsem 1.2.28^73^, prior to normalization on library size with the edgeR^74^ bioconductor package. Finally, differential analysis was conducted with the glm framework likelihood ratio test from edgeR. Multiple hypotheses adjusted p-values were calculated with the Benjamini-Hochberg^75^ procedure to control FDR.

### Single cell RNA sequencing and analysis

For each control and knockout cell line, single cell samples were prepared for 3’ mRNA sequence determinations using the Scipio bioscience protocol (to be published) with barcoding beads (NxSeq Single-cell RNA-seq Beads, LGC Biosearch Technologies). For beads that captured mRNA molecules, reverse transcription, PCR amplification of cDNA, and sequencing library preparation were performed according to published procedures^76^. A total of 35,106,546 reads, 56,108,512 reads, and 54,702,467 reads were generated by NovaSeq from two replicates of *APP-KO2* (KO2-1 and KO2-2) and one CT (control) sample, corresponding to 2000, 2000, and 1500 cells. The reads passed QC process using FASTQC v0.11.8. Sample analyses were performed using UMI-tools v1.0.0. Reads were aligned to GRCh38.94 using STAR v2.7 with default parameters. FeatureCounts v1.6.4^77^ (Ensembl GRCh38.94.GTF) was used to count the number of aligned reads per feature, followed by umi_tools dedup to collapse those reads belonging to a barcode and mapping the same position of a gene. To build the count table umi_tools count was applied. Seurat V3.1.4^39^ was used: i) to filter cells having more than 200 genes and less than 20% mitochondrial genes, with 1326, 1594, and 1299 cells passing the QC filters for the KO2-1, KO2-2, and CT samples, respectively.; ii) to pool both KO samples (2920 total); iii) to cluster KO and CT cells using Seurat clustering with 10 PCA dimensions and resolution 0.5; iv) to identify cluster markers using FindAllMarkers with default parameters. Visualizations were performed with Seurat and ggplot2 packages. DEGs in KO vs CT were analyzed for enrichment in gene ontology, biological and disease associations using Enrichr tools to further explore ChIP-seq datasets linking transcription regulators with DEGs^78^. Integration with other early fetal cells ^22^ was performed using Integrated Anchors analysis followed by clustering with Seurat ^39^.

### Rescue experiments

NPCs of Control and *APP-KO2* were transfected with *pPB-CAG-IRES-EGFP* and *APP-KO2* NPCs were transfected by *pPB-CAG-hAPP-IRES -EGFP* and kept for 4 days. Cells were fixed and stained for GFP/SOX2/DCX. The GFP+ cells were quantified, and cell type was determined by SOX2 and DCX as progenitor and differentiated cell, respectively.

### Neural precursor treatment

WNT3a (R&D Systems) and SR11302 (TOCRIS) were used at final concentrations of 150 ng/ml and 10µM, respectively. Neural progenitor cells of control and *APP-KO2* were treated for 48 hours and then stained for SOX2 and DCX.

### Western blot

Cells were placed on ice, lysed directly in wells by adding RIPA buffer (Sigma) supplemented with Complete protease inhibitor cocktail (Roche), and agitated for 20 min. Thereafter, the samples were collected and centrifuged at >14000 g, 30 min at 4 °C and the supernatant was transferred to a fresh tube. Protein determination was performed using the Pierce™ Detergent Compatible Bradford Assay Kit (ThermoFisher Scientific). Sample buffer (BOLT LDS, Life technologies) was added to equal amounts of protein, and samples were loaded onto a BOLT 4–12% Bis-tris gel and transferred using MES buffer (all from Life technologies). Proteins were blotted onto a 0.2 μM nitrocellulose membrane (GE Healthcare) using semi-dry technique. Membranes were blocked in 5% Nonfat-Dried Milk bovine (Sigma) and incubated overnight at 4 °C with primary antibodies listed in Supplementary Table7. After washing, membranes were incubated with HRP-conjugated secondary antibodies for 1 hour at room temperature. Protein detection was performed by Pierce ECL Western Blotting substrate (ThermoFisher Scientific) and bands were visualized using ChemiDoc™ Touch Imaging System (BioRad Laboratories). Band intensities were calculated using Image Lab software.

### Statistical analysis

Statistical analysis was performed by GraphPad Prism. Data in Fig. panels reflect 3 independent experiments performed on different days. An estimate of variation within each group of data is indicated using standard error of the mean (SEM). We performed unpaired t-test for assessing the significance of differences in the analyses containing two conditions, one-way ANOVA correction in the analyses containing more than three conditions, and two-way ANOVA in the group analysis.

## Acknowledgments

This work was supported by the Investissements d’Avenir program (ANR-10-IAIHU-06), Paris Brain Institute-ICM core funding, a Sorbonne Université Emergence grant, a Neuro-Glia foundation grant and the Roger De Spoelberch Prize (to B.A.H.). Tengyuan Liu is funded by the Chinese Scholarship Council (CSC). We thank Natalia Baumann and Dr. Denis Jabaudon for sharing the pseudotime analysis code, Dr. Bart De Strooper for sharing the anti-APP antibody, Drs. Stephane Nedelec, Joris De Wit, Denis Jabaudon, members of the Hasan lab as well as Antoine Graindorge, and Stuart Edelstein from Scipio biosciences for helpful discussions. Scipio bioscience was supported by the Investissements d’Avenir program and the Région Île-de-France. This work benefited from equipment and services from the ICM’s genotyping and sequencing core facility (iGenSeq). Mouse breeding work was conducted at the PHENO-ICMice facility. We thank Philippe Ravassard for providing the pSYN lentiviral vector backbone and Blandine Bonnamy and Clementine Ripoll from iVECTOR core facility for technical assistance and production of the lentiviral vectors. The Core is supported by 2 Investissements d’Avenir grants (ANR-10-IAIHU-06 and ANR-11-INBS-0011-NeurATRIS) and the “Fondation pour la Recherche Médicale”. iPSC work was carried out at the CELIS core facility with support from Program Investissements d’Avenir (ANR-10-IAIHU-06) Light microscopy was carried out at the ICM.Quant facility. We thank all core technical staff involved especially Stephanie Bigou from CELIS for her advice on iPSCs and CRISPR/Cas9 work and Claire Lovo from ICM.Quant for help with image analysis.

## Author contribution

Kh.Sh., C.P., and B.H. conceived the project, interpreted the data, and wrote the manuscript with input from all authors. Kh.Sh. and B.H. designed, and Kh.Sh. performed experiments and data analysis for most of the project. J.P. performed experiment on NPCs derived from the second iPSC line and 3D culture derived from both iPSC lines. M.B.T.Z performed the viral approach experiment. T.L performed mouse embryo work. J.K. performed sample preparation for single cell RNA-seq A.S. and C.P performed single cell data analysis. N.D. generated the vectors for the rescue experiment. E.L. and D.B. performed motor neuron culture and staining. R.L. helped optimize the iPSC and NPC culture and differentiation protocols.

